# Effects of Aging on the Immune and Periosteal Response to Fracture in Mice

**DOI:** 10.1101/2024.11.06.622348

**Authors:** Justin S King, Matthew Wan, Adam Kim, Sanja Novak, Shagun Prabhu, Ivo Kalajzic, Anne M Delany, Archana Sanjay

## Abstract

Aging predisposes individuals to reduced bone mass and fragility fractures, which are costly and linked to high mortality. Understanding how aging affects fracture healing is essential for developing therapies to enhance bone regeneration in older adults. During the inflammatory phase of fracture healing, immune cells are recruited to the injury site as periosteal skeletal stem/progenitor cells (pSSPCs) rapidly proliferate and differentiate into osteochondral lineages, allowing for fibrocartilaginous callus formation and complete bone healing. Irrespective of age, how periosteal mesenchymal and immune cells interact during early fracture healing is incompletely understood, limiting our ability to potentially modulate these processes. To address this, we directly analyzed, in parallel, at a single-cell level, isolated murine CD45(+) and CD45(-) periosteal cells dissected from intact and fractured bones, collected three days after injury. Through comprehensive analysis, corroborated by bulk RNA-sequencing, flow cytometry, and histology, we found aging decreases pSSPCs proliferative, marked by a reduced expression of genes required for callus formation and an increased senescence signature. We found that the chemokine *Cxcl9* was highly upregulated in aged intact Prrx1+ pSSPCs, predicted to interact with other pSSPCs directly, and associated with increased recruitment of CD8+ T cells at the fracture site three days after injury. Cell-to-cell communication analysis provided insight into the complexity of interactions among the many cell types regulating fracture healing and the impact of aging on these processes. Together, these results provide insight into age-induced alterations in fracture healing, informing the development of improved therapeutic approaches for fragility fractures.

## Introduction

Aging heightens the risk of osteoporosis, fracture susceptibility, and impedes regeneration, presenting an emerging public health concern in the face of an aging population (Goodnough & Goodman, 2022; Lewiecki et al., 2019; López-Otín et al., 2023). This is exemplified by fragility fracture of the hip associated with a one-year mortality rate approaching 30%, with projected healthcare costs for fragility fractures expected to increase by 70% from 2018 to 2040 (Brauer, 2009; Lewiecki et al., 2019). A better understanding of the age-related changes in skeletal resilience and fracture healing is needed to improve treatment strategies and develop new therapies to address these significant socioeconomic costs.

Skeletal stem/progenitor cells (SSPCs) are crucial for skeletal regeneration, functioning as the source of matrix-producing cells for bone maintenance and healing following injury. While aging affects SSPCs’ proliferation and differentiation potential, these cells are inherently heterogeneous. Distinct SSPC subtypes exist across different skeletal regions, such as the skull, vertebrae, and long bones, with further variability within the long bones themselves; distinct SSPC subtypes have been identified within the diaphysis, metaphysis, bone marrow (bmSSPCs), and periosteal compartments (pSSPCs) (Bok et al., 2023; Chan et al., 2015; Debnath et al., 2018; Li et al., 2022; Sun et al., 2023; Yang et al., 2024). Periosteal SSPCs are essential for intramembranous bone formation, and their role in forming new bone through endochondral ossification of a fibrocartilage callus during fracture healing illustrates their plasticity. In aged mice, reduced expression of cartilaginous genes during early healing results in a smaller fibrocartilage callus volume and delayed healing (Bahney et al., 2019; Matthews et al., 2014). However, these studies did not examine the contribution of the periosteal response, despite the periosteum being the principal source of cells forming the fibrocartilage callus.

Immune cells also play a critical role in bone repair, particularly during the inflammatory phase occurring in the first few days after fracture. Neutrophils and monocytes/macrophages rapidly infiltrate the fracture site to clear debris and initiate healing. Emerging evidence suggests that other immune cell subtypes, including mast cells and T cells, are also crucial for coordinating the repair process (Baht et al., 2018; Kroner et al., 2017). In aging, there is an altered immune response, with increased systemic inflammation that may interfere with SSPC function, ultimately affecting the efficiency of bone regeneration (Goodnough & Goodman, 2022; Josephson et al., 2019; Molitoris, Huang, et al., 2024).

Most research on how aging impacts fracture healing has focused on bone marrow and endosteal cells, revealing that aging alters SSPC transcriptional profiles that drive osteoclastic activity, reduces SSPC diversity and number, and increases cellular senescence (Ambrosi et al., 2021; Josephson et al., 2019; Remark et al., 2023). These age-related changes collectively lead to diminished fracture healing efficiency; however, less is known about how aging impacts the periosteum. To examine how aging affects the periosteum and its acute response to fracture, we directly analyzed, in parallel, at a single-cell level, isolated CD45(+) and CD45(-) periosteal cells dissected from young and old mice, both intact and three days after fracture – a critical period when pSSPCs are rapidly dividing and beginning to differentiate into osteochondral lineages (Novak et al., 2024). Our findings show a decline in the proliferation capacity of pSSPCs, increased cellular senescence, and decreased expression of chondrogenic genes in aged mice, alongside an increase in CD8+ cells at the fracture site. Further, we predicted cell-to-cell interaction networks between the mesenchymal, endothelial, and immune cells during early fracture. These results could help inform regenerative strategies to promote bone healing, and our uniquely comprehensive and matched scRNA-seq datasets will serve as a resource for the broader orthopedic research community.

## Results

### Periosteal cell proliferation in response to fracture is decreased in aged mice

While bmSSPCs are the primary contributing cell type to bone remodeling and the intramembranous ossification and healing of completely stabilized bone injuries, pSSPCs give rise to most of the mesenchymal cells that contribute to the fibrocartilaginous callus and regenerated bone during healing of partially stabilized fractures (Jeffery et al., 2022; Perrin & Colnot, 2022). In the first few days after a fracture, pSSPCs rapidly divide and differentiate into osteochondral lineages, initiating the healing process that sets the stage for complete repair (Julien et al., 2022; Y. L. Liu et al., 2024; Matthews et al., 2014; Novak et al., 2024). To study how aging affects the periosteal response to fracture, we assessed differences in the cellular responses between young (∼3 months) and aged (20-24 months) mice that underwent a partially stabilized transverse femoral fracture. Harvesting the femurs that were either intact or three days post-fracture, we assessed cell proliferation and early osteochondral differentiation (Fig 1A-C). To identify cycling cells, indictive of cell proliferation, we injected EdU ((5-ethynyl 2’-deoxyuridine) 24 hours before tissue harvest and observed that three days post-fracture, young mice significantly increased EdU+ cells compared to aged mice (P<0.0001). In contrast, aged mice showed a less pronounced increase in EdU+ cells (P=0.0555) (Fig 1B-C).

**Figure 1.**
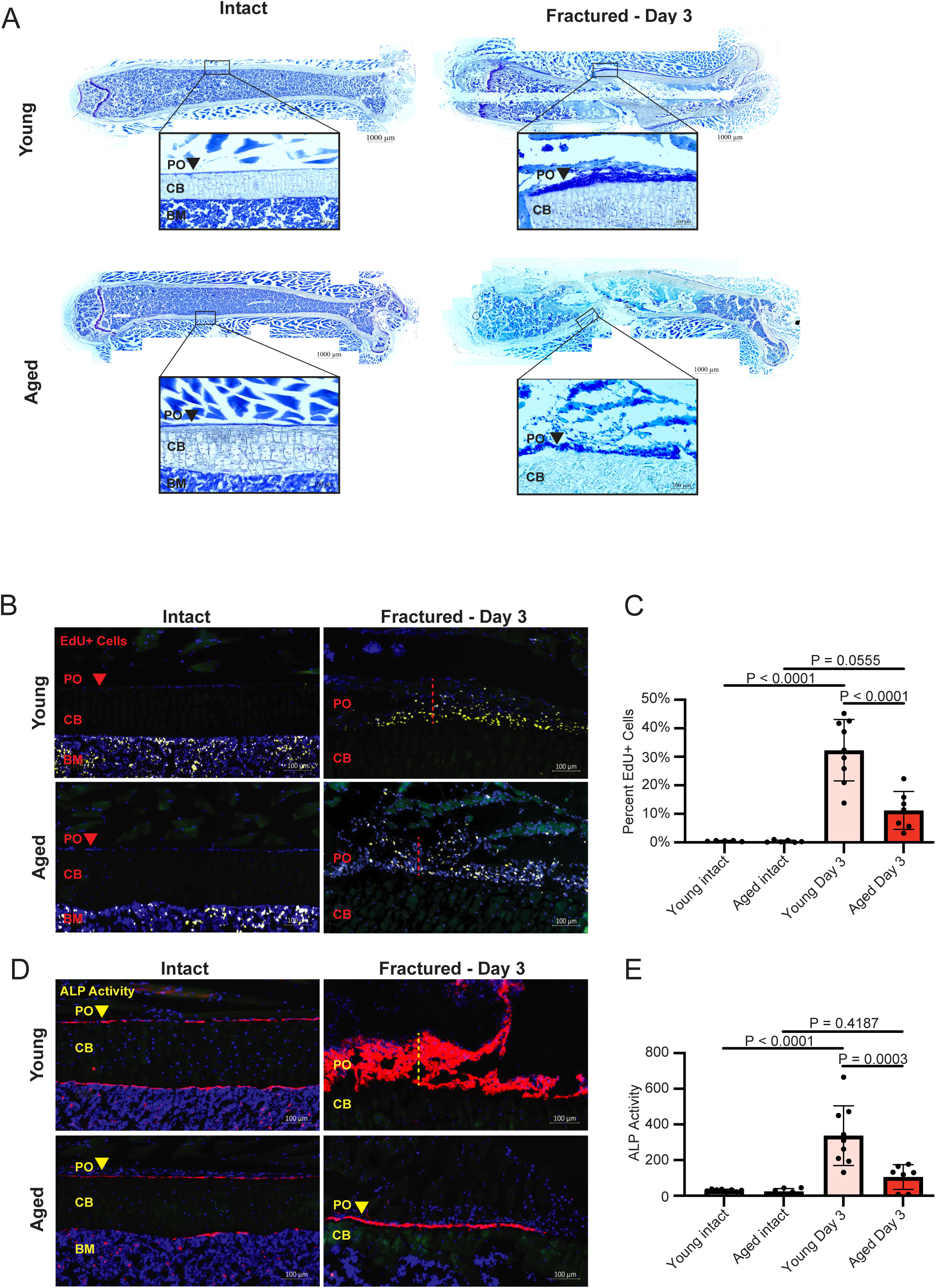
Stitched tile scan images and high-magnification images of representative cryosections from intact and day three post-fracture femurs stained with Toluidine blue, collected from young and aged mice; zoomed-in portions indicate the regions from which images in panels B and D were taken from adjacent cryosections. **B**) Cryosections of intact femurs and femurs collected three days post-fracture, with EdU injected 24 hours prior to tissue harvest to label proliferating cells. **C)** Quantification of EdU-positive cells. **D)** Alkaline phosphatase (ALP) activity staining to assess osteogenic activity. **E)** Quantification of ALP activity, represented as pixel intensity normalized to area. Abbreviations: PO = periosteum; CB = cortical bone; BM = bone marrow. Dashed lines indicate the thickened periosteum following fracture. All images were taken from cryosections, with adjacent sections from the same bone used for each condition. Representative images are shown; n = 5-9 mice per condition. Statistical analysis was conducted using a one-way ANOVA test followed by Tukey’s multiple comparisons test.

Additionally, we assessed alkaline phosphatase (ALP) activity, an early marker of osteochondral differentiation potential, at the same time points. Young mice exhibited a robust increase in ALP activity post-fracture (P<0.0001), whereas aged mice showed no significant change in ALP activity (P=0.4187) (Fig 1D-E). Together, these findings indicate that pSSPC proliferation is reduced with aging, marked by lower EdU incorporation, resulting in decreased cellularity. This reduction in cellularity also likely contributes to the diminished ALP activity.

### Assessing the periosteum and periosteal response to fracture injury at the single-cell level

To better understand the impact of aging on the cellular dynamics of fracture healing, we performed droplet-based single-cell RNA sequencing (scRNA-seq) on femoral periosteal cells. Our groups included cells harvested from young and aged mice, intact and three days post-fracture, pooling cells from 5-7 mice per group (Fig 2A). To ensure that our results better reflect native cellular states without artifacts introduced by *ex vivo* manipulations, we did not culture or expand the cells before sequencing (Zhang et al., 2023). To enrich for non-hematopoietic mesenchymal cell lineages and to analyze interactions between cell types, we FACS-sorted Ter119(-)CD45(-) and Ter119(-)CD45(+) cells to perform scRNA-seq on both groups in parallel, loading 6,000 cells, in equal number, for each group (Fig 2B). In doing so, our FACS indicated that CD45(-) cells expanded from 2.9% to 7.9% in young mice and from 2.2% to 3.5% in aged mice, consistent with the reduced cell expansion observed in our EdU data three days post-fracture (Fig 1B).

**Figure 2.**
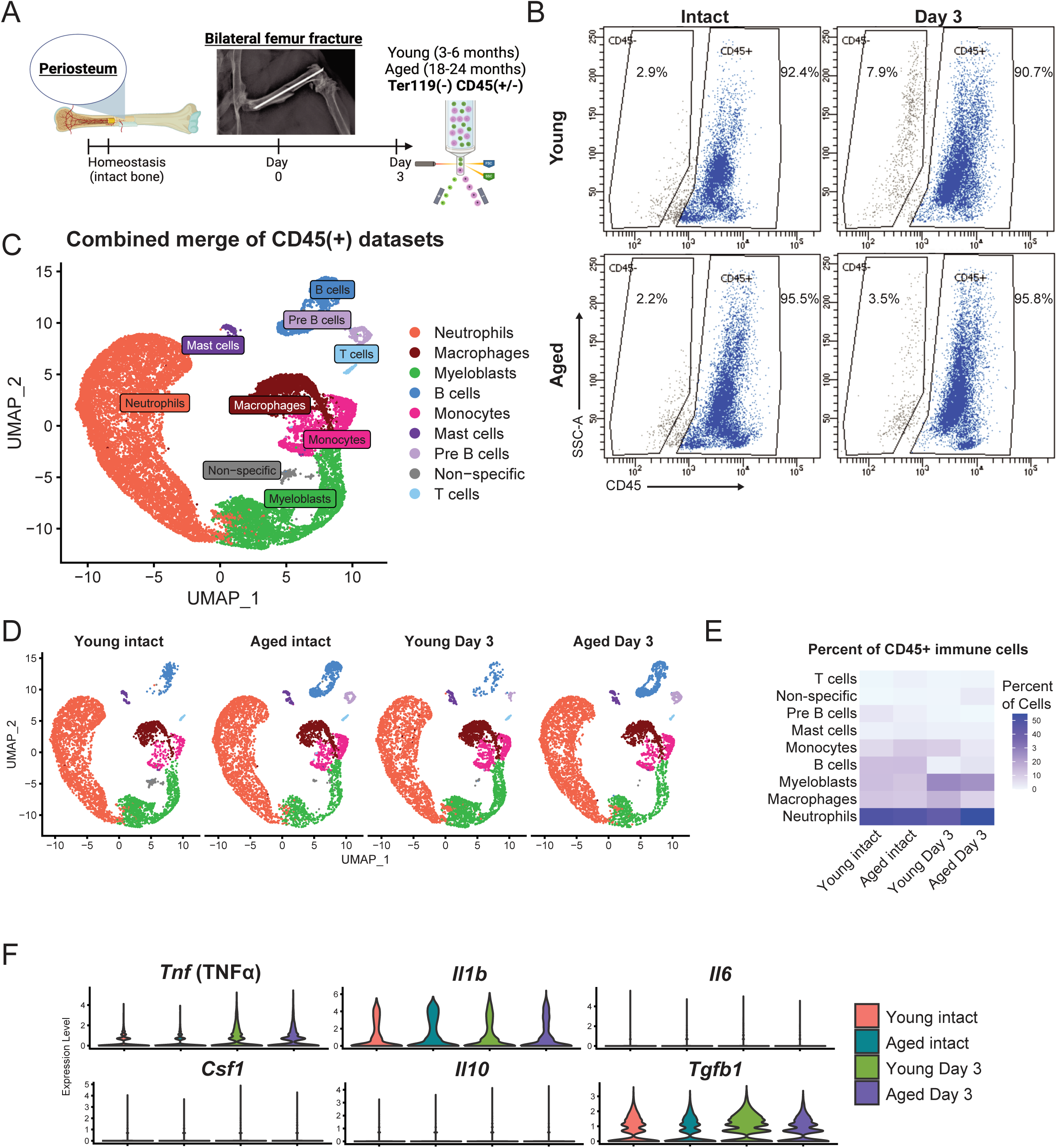
Periosteal response to fracture at the single-cell level. **A)** Schematic of the experimental approach to obtain periosteal cells from intact and three days post-fractured femurs by fluorescence-activated cell sorting (FACS) for equal numbers of CD45(+) and CD45(-) cell populations prior to scRNA-sequencing. **B)** FACS plots after gating for Ter119(-) cells, showing the percentage of CD45(-) or CD45(+) cells from the parent population; these gated populations were used to collect CD45(-) and CD45(+) for scRNA-seq **C)** Initial merged and annotated clustering of CD45(+) scRNA-seq datasets, **D)** which were then split apart by experimental group. **E)** Heatmap of immune cell types identified by scRNA-seq, plotted as percent of cell type identified in each condition. **F)** Violin plots for cytokines implicated in fracture healing and bone regeneration from all cells in each CD45(+) scRNA-seq condition.

### Profiling immune cells within the periosteum by scRNA-seq

We began by analyzing the scRNA-seq data from the CD45(+) datasets based on marker gene expression. (Fig 2C; S1A-C). Although transcripts associated with osteoclasts (*Acp5*, *Ocstamp*) were present in the monocyte and macrophage clusters, we could not identify a specific cluster for osteoclast precursors or mature osteoclasts. Additionally, there was a significant population of myeloblasts (*Cd34*) (Fig 2D-E). After fracture, the populations of cells identified as macrophages and myeloblasts increased in the periosteum, particularly in young mice (Fig 2E).

### Cytokine expression within periosteal immune cells

Next, we analyzed cytokine and chemokine gene expression within our CD45(+) scRNA-seq datasets to understand potential changes during fracture repair (Fig 2F; Fig S2) (Bahney et al., 2019). In murine fracture models, TNF-α, IL-1β, and IL-6 are typically expressed at the injury site within 24 hours of injury (Cho et al., 2002; Kon et al., 2001). The expression of TNF-α in fractures follows a biphasic pattern, peaking initially during the onset of fracture repair and again during the transition from chondrogenesis to osteogenesis in endochondral maturation (Kon et al., 2001; Lehmann et al., 2005). Consistent with the role of TNF-α in modulating fracture healing through multiple pathways (Gerstenfeld et al., 2003; Lehmann et al., 2005; Molitoris, Huang, et al., 2024), we observed an increase in *Tnf* expression in CD45(+) cells, following fractures in both young and aged mice, although the differences between the age groups were not significant. The TNF was largely sourced from the macrophage population (Fig 6D). IL-1β (*Il1b*) expression was more robust in intact aged mice, aligning with literature that associates increased IL-1β with aging (Sayed et al., 2021). After fracture, there were no significant differences in *Il1b* between the age groups. We noted low expression of IL-6 (*Il6*) and IL-10 (*Il10*), with mast cells predominantly expressing *Il6*, *Il4*, *Il10*, and *Csf1*, which are cytokines not significantly expressed by other immune cell populations (Fig S2). These observations align with reports that IL-6-deficient mice do not exhibit significant impairment in healing outcomes when subjected to stress or transverse fracture models (26–28). Upregulation of TGF-β1 (Tgfb1) was also observed in CD45(+) periosteal cells after fracture, which was reduced in cells from aged mice (Fig 2F).

### Assessment of CD45(-) populations in the intact and injured periosteum from young and aged mice

Within our CD45(-) data set (Fig 3A), we assessed markers previously identified in lineage-tracing experiments to characterize pSSPCs that give rise to matrix-producing cells. These pSSPC clusters were marked by Paired related homeobox 1 (*Prrx1*) (Jeong et al., 2024), Cathepsin K (*Ctsk*) (Debnath et al., 2018), α-smooth muscle actin (αSMA; *Acta2*) (Matthews et al., 2021), *Gli1* (Jeffery et al., 2022; Shi et al., 2017), and Platelet-derived growth factor receptor alpha (*Pdgfra*) (Xu et al., 2022), Leptin receptor (*Lepr*), Adiponectin (*Adipoq*), (Jeffery et al., 2022), CD51 (*Itgav*), and CD90 (*Thy1*) (Chan et al., 2015)(Fig 3A). Within the CD45(-) dataset, we also identified clusters of endothelial cells, myocytes/muscle, mature osteochondral cells, erythroblasts, and residual hematopoietic cells (Fig 3B; Fig S3B).

**Figure 3.**
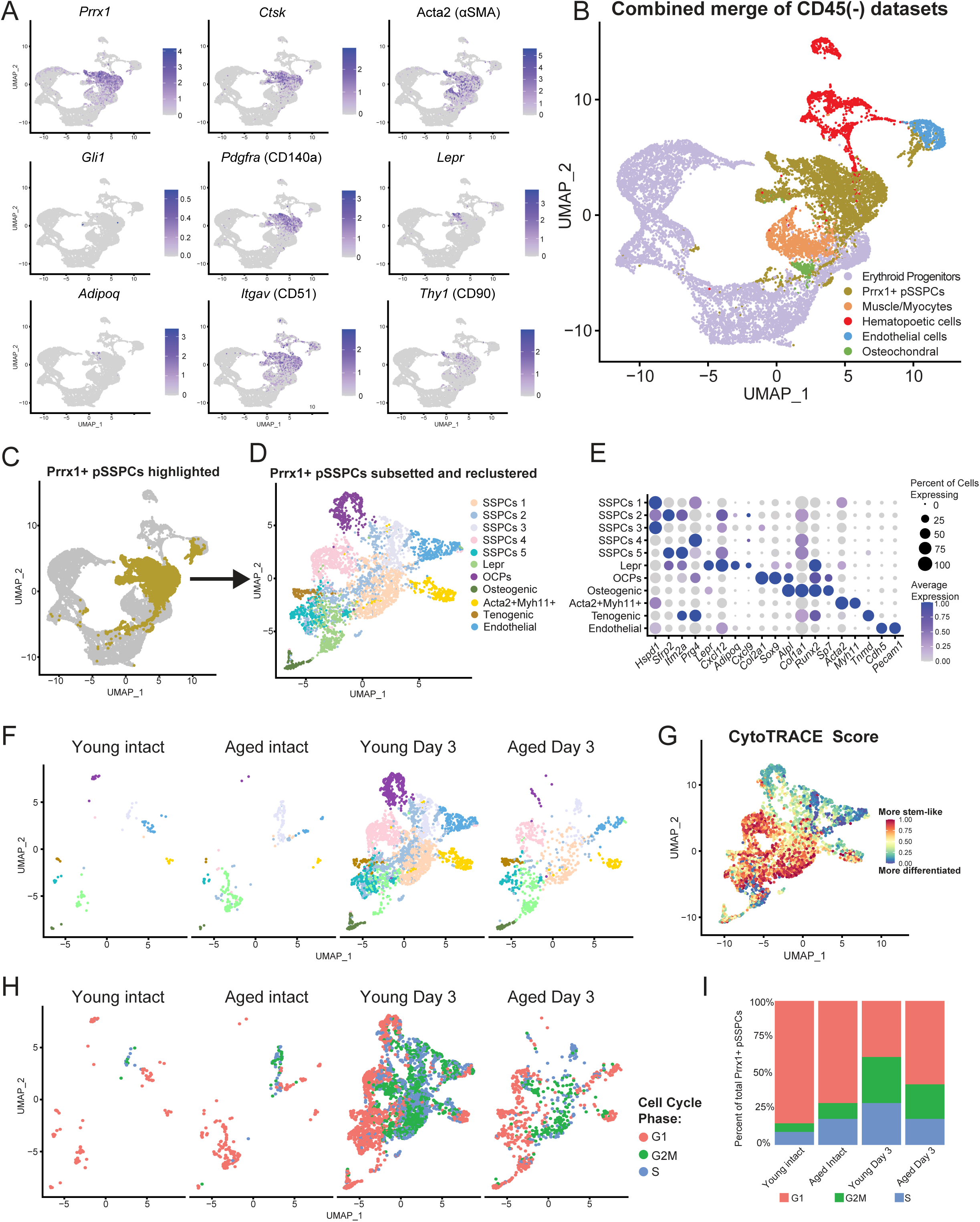
Analysis of CD45(-) mesenchymal cells from the periosteum at the single-cell level. **A)** Expression pattern of selected periosteal SSPC markers. **B)** Initial clustering of cell types identified in merged scRNA-seq datasets isolated from CD45(-) periosteal cells from all groups. **C)** The Prrx1+ pSSPC cluster is highlighted in gold was **D)** subsetted and reclustered. **E)** Marker genes for new clusters identified the Prrx1+ pSSPCs population. **F)** Merged subsetted and reclustered Prrx1+ pSSPC population in each experimental group. **G)** In merged Prrx1+ pSSPCs, the CytoTRACE score is assigned to each cell. **H)** Prrx1+ pSSPCs in each experimental group bioinformatically assigned a cell-cycle state, and **I)** quantified.

As *Prrx1* was the most broadly expressed among the previously known putative pSSPC markers examined and likely represented the subset of cells that facilitate the formation of the fibrocartilage callus and new osteoblasts, we subsetted and reclustered these cells as ’Prrx1+ pSSPCs’ (Fig 3C-D; Fig S3C). Distinct clusters within these cells included those reprehensive of tenogenic cells (*Tnmd*), osteogenic cells, osteochondral precursors (OCPs; Col2a1, Runx2), *Lepr*+ cluster, pericyte cells (aSMA+Myh11+; *Acta2, Myh11*), residual endothelial cells (*Chd5, Pecam1)*, and several clusters of pSSPCs, annotated as SSPC 1-5 (Fig 3D-E). The SSPC 1-5 clusters were enriched with marker genes associated with periosteal, chondrogenic, and osteogenic regenerative processes in the skeleton, including *Itm2a* (Xing et al., 2024), *Sfrp2* (De Castro et al., 2021), *Prg4* (Krawetz et al., 2022) and *Hspd1* (Suwanwela et al., 2011) (Fig 3E). The Prrx1+ SSPCs constituted 4.0% and 6.3% of the CD45(-) cells in intact young and aged mice, respectively. After a fracture, the proportion of Prrx1+ pSSPCs increased to 54.5% in young and 28.9% in aged mice, representing a 13-fold and 4.5-fold increase, respectively (Fig 3F). The ’SSPC 1’ cluster expanded the most after fracture and also had the highest “stem-like” CytoTRACE score, a metric used to infer stemness and differentiation state based on the transcriptome (Gulati et al., 2020) (Fig 3G; Fig S3D-E).

### Cell cycle changes in the periosteum with age

To further investigate the cause of decreased cell expansion among Prrx1+ pSSPCs, we performed an analysis of cell cycle genes and bioinformatically assigned a cell cycle state to these cells (Fig 3H). We found that most cells from both intact young and aged mice were in the non-cycling G0 or G1 phase. However, 14.5% of Prrx1+ pSSPCs in uninjured young mice were in the G2M or S phase, which increased to 60.6% post-injury. In contrast, among Prrx1+ pSSPCs in aged mice, 29.3% were cycling in uninjured mice, while only 42.5% were cycling post-injury (Fig 3I). The impaired expansion of Prrx1+ pSSPCs in aged mice is consistent with our EdU and FACS data (Fig 1B, C; Fig 2B), reinforcing age-related changes in periosteal cell dynamics in response to injury.

### Cxcl9 expression increases in the periosteum with age

We then examined differential gene expression within Prrx1+ pSSPCs, comparing the intact young and aged groups, assessing for factors that may compromise fracture healing before the process begins. The chemokine *Cxcl9* (C-X-C motif chemokine ligand 9) was one of the most differentially upregulated genes in Prrx1+ pSSPCs derived from aged mice, particularly expressed in the SSPC 2, SSPC 4 and Lepr+ populations (Fig 3B; Fig. 4A-C; Fig S4A). *Cxcl9* was also upregulated in aged intact CD45(+) periosteal cells, particularly within the monocyte and macrophage populations (Fig S4B). While CXCL9 is known to regulate T-cell function via binding to its receptor CXCR3 (Chen et al., 2019), *Cxcr3* is minimally expressed in our CD45(+) scRNA-seq dataset and was not detected in our CD45(-) dataset (Fig S4A-B). However, through CellChat analysis of our integrated CD45(+) and CD45(-) scRNA-seq datasets, SSPC-derived *Cxcl9* was predicted to interact with the Atypical chemokine receptor, encoded by the *Ackr1* gene (Fig 4D). In young intact periosteum, Cxcl9 synthesized by SSPC 2 cells is predicted to interact primarily with Ackr1 on SSPC 5, whereas in aged intact periosteum Cxcl9 synthesized by SSPC 2, SSPC 3, and Lepr+ cells were predicted to interact with Ackr1 on SSPC 5 population. While the Cxcl9 – Cxcr3 interaction is within the CellChat ligand – receptor database, no interaction between Cxcl9 and Cxcr3 was detected during the analysis of our data (Fig 3E).

**Figure 4.**
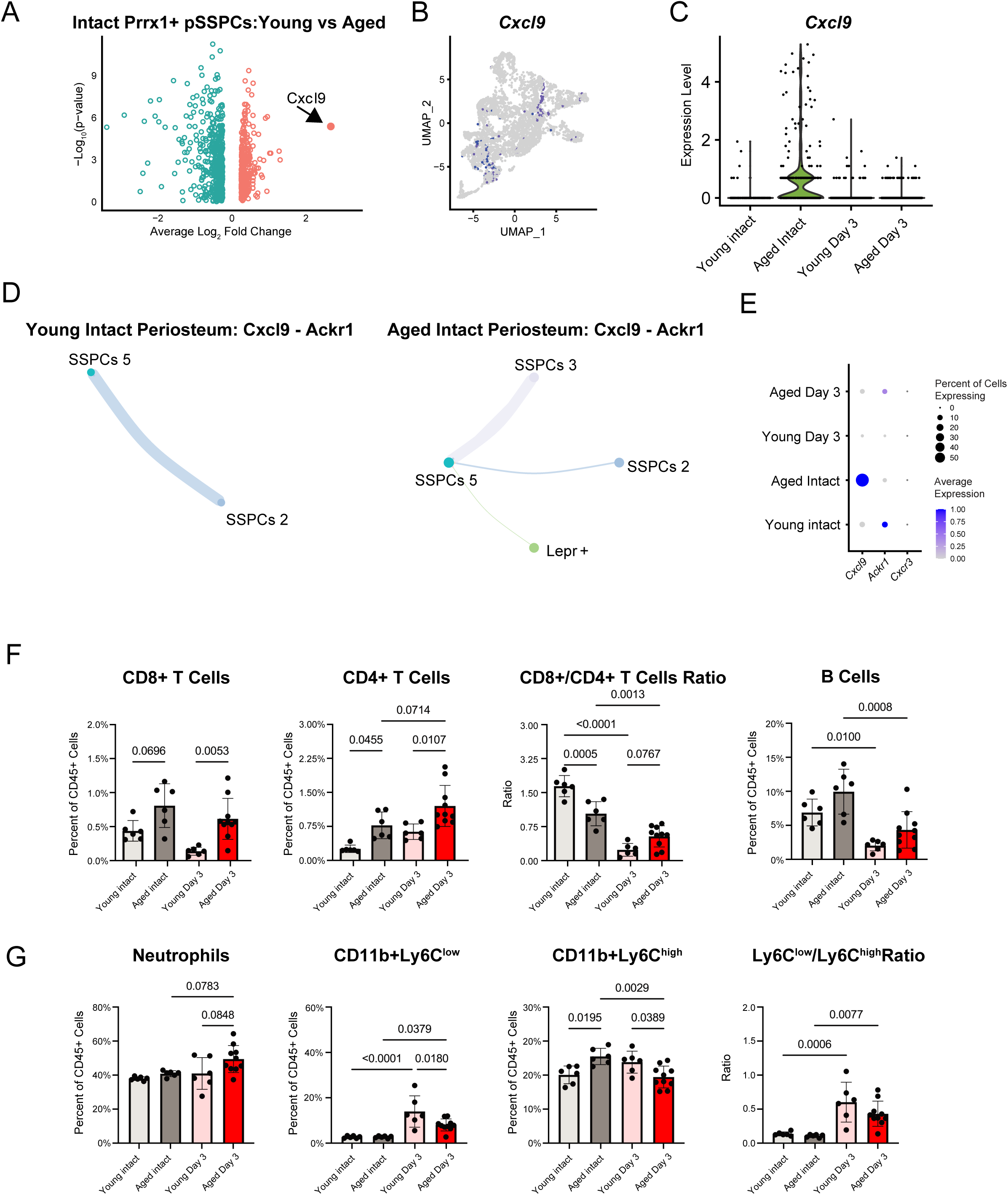
Cxcl9 is upregulated Prrx1+ pSSPCs from aged intact mice. **A)** Volcano plot of differential gene expression comparing intact young and intact aged Prrx1+ pSSPCs . **B)** Feature plot of expression pattern of *Cxcl9* in merged Prrx1+ pSSPCs scRNA-seq data. **C)** Violin plot showing the expression of *Cxcl9* in the subsetted Prrx1+ pSSPCs for each group. **D)** CellChat Analysis showing Cxcl9 – Ackr1 interaction obtained using merged annotated datasets from both the CD45(-) and CD45(+) scRNA-seq groups; the color of the line corresponds to the color of the dot representing the “sending cell” group within the interaction; a thicker line represents a stronger interaction. **E)** Dot plot for *Cxcl9*, *Ackr1*, and *Cxcr3* expression in Prrx1+ pSSPCs. **F)** Flow cytometric analysis of periosteal lymphocytes and **G)** myeloid-lineage immune cells; each dot represents one biological replicate; n=6-10 male mice/age/timepoint. Statistics were performed using an ordinary One-Way ANOVA followed by Tukey’s multiple comparisons. P-values reported for all values <0.10.

### Profiling immune cells within the periosteum by flow cytometry

CXCL9 is a CD8+ T cell chemoattractant (Chen et al., 2019), and since immune dysfunction and inflammation are known to contribute to impaired fracture healing, we assessed discrete immune cell populations in the periosteum by flow cytometry (n=6-10) (Fig 4F; antibodies detailed in Table S2; dot plots and gating strategy in Fig S5). First, we analyzed CD3+CD4+ (helper) and CD3+CD8+ (cytotoxic) T cell populations. The percentage of CD8+ T cells increased significantly post-fracture in aged mice compared with young (P=0.0053). Among CD4+ T helper cells, aged mice exhibited a higher proportion in intact bone compared to young (P=0.0455), with this difference becoming more pronounced after fracture (P=0.0107). Overall, the CD8+/CD4+ ratio was significantly higher in intact bone compared to post-fracture in both age groups (P<0.01 in both cases; Fig 4F). In contrast, there were no significant differences in the percentage of B cells (CD45+B220+) between young and aged mice. However, the B cell proportion decreased three days post-fracture in both young (P=0.0100) and aged (P=0.0008) mice, potentially due to increased recruitment of other hematopoietic cells.

Among myeloid lineage cells (Fig 4G), the increase in percentage of neutrophils (CD45+CD11b+Ly6G+) in periosteal tissues post-fracture in aged mice compared with young did not reach statistical significance (P=0.0848). The monocyte/macrophage population was subdivided into ‘patrolling’ (CD11b+Ly6C^low^; M2-like) and ‘inflammatory’ (CD11b+Ly6C^high^; M1-like) subsets for analysis (Kou et al., 2024; Molitoris, Balu, et al., 2024). Young mice showed a significant increase in patrolling monocytes post-fracture (P<0.0001), while this increase was significantly attenuated in aged mice (P=0.0180). In contrast, inflammatory monocytes were elevated in aged intact periosteum compared with young (P=0.0195), and these cells decreased markedly in aged mice post-fracture (P=0.0029), such that they were lower than the levels observed in young mice after fracture (P=0.0389). In contrast, the percentage CD11b+Ly6C^high^ inflammatory monocytes in young mice was similar in both intact periosteum and at three days post-fracture. Consequently, the Ly6C^low^/Ly6C^high^ monocyte ratio significantly increased three days post-fracture in both young (P=0.006) and aged (P=0.0077) mice, suggesting an infiltration of patrolling macrophages into the periosteum. As the periosteum is heavily vascularized, it is possible that changes in immune cells present in the periosteum may be attributed to changes to immune cells found in the greater vasculature and blood. To test this hypothesis, we also assessed the immune cell composition in the peripheral blood obtained by cardiac puncture and in bone marrow flushed from the same mice used for periosteal cell harvesting. Our results found large variations in the immune cell distribution between the peripheral blood, bone marrow, and periosteum, suggesting that each compartment has a unique immune cell composition (Fig S6).

### Osteochondral changes during the periosteal response to fracture

To assess how aging affects gene expression three days after fracture, we analyzed all the differentially expressed genes within Prrx1+ pSSPCs post-fracture (Fig 5A). Genes associated with chondrogenesis, such as *Col2a1* and *Acan*, were among the most enriched in young adult mice but were downregulated with aging. Aged mice also exhibited downregulation of Periostin (*Postn*) and upregulation of *Csf1* post-fracture, both markers associated with age-related impairments in fracture healing (Clark et al., 2023) (Ambrosi et al., 2021). To investigate cellular processes involved in the initiation of bone repair, we conducted Gene Set Enrichment Analysis (GSEA) using the Gene Ontology Biological (The Gene Ontology Consortium et al., 2023) gene sets on differentially expressed genes in Prrx1+ pSSPCs from young versus aged mice at three days post-fracture. The analysis revealed that gene sets related to chondrogenic processes and matrix deposition were predominantly enriched in young mice. In contrast, gene sets enriched in aged mice included ’cell killing’ and ’defense/immune responses’, which include many genes involved in inflammation, suggesting that inflammaging may contribute to decreased bone healing efficacy with age (Fig 5B) (Josephson et al., 2019).

**Figure 5.**
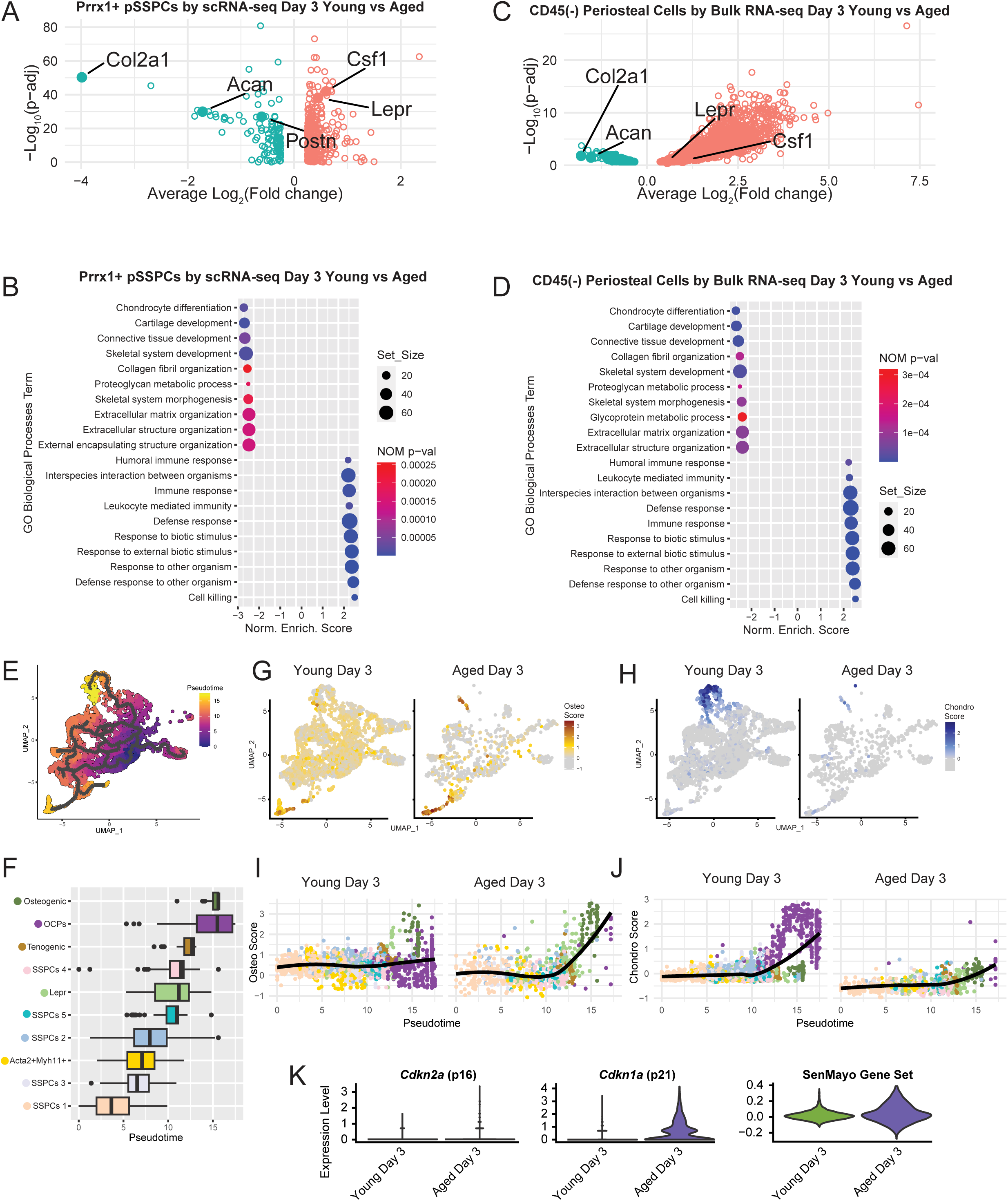
Chondrogenic gene expression decreases three days post-fracture in the periosteum of aged mice. **A)** Volcano plot of differentially regulated genes among scRNA-seq identified Prxx1+ pSSPCs from periosteum three days post-fracture of young versus aged mice. **B)** Top 20 upregulated and downregulated Gene Ontology (GO) Biological Processes identified by scRNA-seq of Prxx1+ pSSPCs three days post-fracture, comparing young versus aged. **C)** Volcano plot of differentially regulated genes identified from bulk RNA-sequencing of CD45(-) isolated cells collected from periosteum three days post-fracture, comparing young versus aged mice. **B)** Top 20 upregulated and downregulated Gene Ontology (GO) Biological Processes identified by bulk RNA-sequencing of CD45(-) isolated cells collected from periosteum three days post-fracture of young versus aged mice. **E)** Monocle 3 trajectory analysis of Prrx1+ pSSPCs, with endothelial cells removed, which **F)** yielded the *Pseudotime* for each cluster. **G)** *Osteo Score* and **H)** *Chondro Score* of reclustered Prrx1+ pSSPCs identified from scRNA-seq analysis of periosteum from young and aged mice three post-fracture. **I)** The *Osteo Score* and **J)** *Chondro Score* of each cell in the Prrx1+ pSSPC cluster plotted against *Pseudotime*. **K)** Senescence burden among Prxx1+ pSSPCs from young and aged mice three post-fracture estimated using the SenMayo gene set.

To validate our single-cell RNA sequencing (scRNA-seq) findings, we performed bulk RNA-seq on CD45(-) cells isolated from the periosteum of a separate cohort of young and aged mice three days post-fracture (n=3-5) (Fig S7A). The differentially expressed genes identified in our bulk RNA-seq dataset were consistent with those from our scRNA-seq analysis, including downregulation of *Col2a1* and *Acan* and upregulation of *Lepr* and *Csf1* in aged mice. Similarly, GSEA using Gene Ontology Biological Process (GO BP) on the differentially expressed genes identified through bulk RNA-seq data showed almost congruent results with our scRNA-seq GSEA (Fig 5C-D). These findings indicate that single-cell and bulk RNA sequencing approaches consistently and robustly identified differential regulation of chondrogenic gene programs between young and aged mice three days post-fracture.

Since the chondrogenic plasticity of periosteal stem cells is essential for proper fracture healing (Debnath et al., 2018), the downregulation of genes associated with chondrogenesis suggests impairments in pSSPC differentiation. To investigate this hypothesis, we used Monocle 3 to create a trajectory to generate cell state *Pseudotime* for the Prrx1+ pSSPCs. For this assessment, we removed the residual endothelial cells and used the ’SSPC 1’ cluster as the starting point in the trajectory (Fig 5E-F) because the ’SSPC 1’ cluster showed the highest degree of stem-ness as determined by CytoTRACE (Fig 3G; Fig S3D-E). Mapping the *Pseudotime* to each cluster revealed that tenogenic, osteochondroprogenitor (OCPs), and osteogenic cells were the most differentiated cell clusters. By assigning each cell an *Osteo Score* or *Chondro Score* based on the aggregate expression of genes implicated in osteogenesis and chondrogenesis (Table S1), we were able to better gauge the assumed identity of the pSSPCs in the periosteal response to injury (Fig 5G-H). Mapping the Pseudotime and Osteo Score or Chondro Score for each cell showed a dramatic decrease in the chondrogenic signature in aged pSSPCs, while the osteogenic signature was elevated in aged pSSPCs (Fig 5I-J). These findings suggest that the fracture-induced chondrogenic plasticity required for endochondral ossification and initial cartilaginous callus formation by periosteal stem cells (Debnath et al., 2018) is impaired with aging.

### Cellular senescence in periosteal cells

As studies have repeatedly demonstrated that cellular senescence negatively affects skeletal resiliency (Doolittle et al., 2023; J. Liu et al., 2022; Saul & Khosla, 2022), we assessed p16 (*Cdkn2a*), p21 (*Cdkn1a*), and markers of the Senescence-Associated Secretory Phenotype (SASP), a recognized hallmark of aging characterized by the secretion of pro-inflammatory factors and matrix metalloproteases by senescent cells (López-Otín et al., 2023). To assess SASP, we used the SenMayo Gene Set, which includes dozens of genes associated with senescence, extending beyond p16 and p21 (Fig 5K; Table S1) (Saul et al., 2022). In aged mice at three days post-fracture, p16 expression showed only a minimal increase compared with young, while p21 expression was dramatically elevated, consistent with the literature (Saul et al., 2024). Additionally, the SenMayo Gene Set composite score was higher in aged mice both before and after fracture (Fig 5K; Fig S7B). Furthermore, the elevated SenMayo score was predominantly localized to the mesenchymal and endothelial cells in our CD45(-) dataset (Fig S7C). In contrast, there was less variation in p16 and p21 expression between age groups in our CD45(+) scRNA-seq datasets (Fig S7D).

### Cell-to-cell communication networks between CD45(+) and CD45(-) populations

To better understand cell-to-cell communication networks at homeostasis and during fracture healing, we integrated our CD45(+) and CD45(-) datasets for each condition and performed a CellChat analysis (Jin et al., 2021). For each time point and condition, we integrated data from annotated immune cell types from our CD45(+) datasets, Prrx1+ pSSPC subclusters, and other cell populations such as endothelial cells, muscle, and mature osteochondral cells from the CD45(-) dataset. Since *Csf1* expression was upregulated with aging in Prrx1+ pSSPCs (Fig 6A), and previous studies have reported that age-induced overexpression of CSF1 in skeletal stem cells contributes to age-related inefficiencies in fracture healing (Ambrosi et al., 2021), we examined the interaction between *Csf1* and its receptor, *Csf1r*. The analysis showed that Csf1 is predominantly derived from the ’Lepr’ and ’SSPC’ clusters in intact periosteum, with its activity focused on the monocyte and macrophage clusters, consistent with the mechanism of CSF1 in the bone marrow as a growth factor for monocytes and macrophages (Inoue et al., 2023). The analysis revealed that mast cells are also a major source of *Csf1*, especially after fracture in young mice. We identified several key pathways that were significantly upregulated in aged samples, including IL-10 signaling, classically associated with anti-inflammatory processes, which was more prevalent in young mice (Fig 6B), and pathways classically associated with pro-inflammatory processes such as *Tnf* (TNFα) and its interaction with its receptor Tnfrsf1a (TNF Receptor 1), which were more prevalent in aged mice (Fig 6C)(Maruyama et al., 2020).

**Figure 6.**
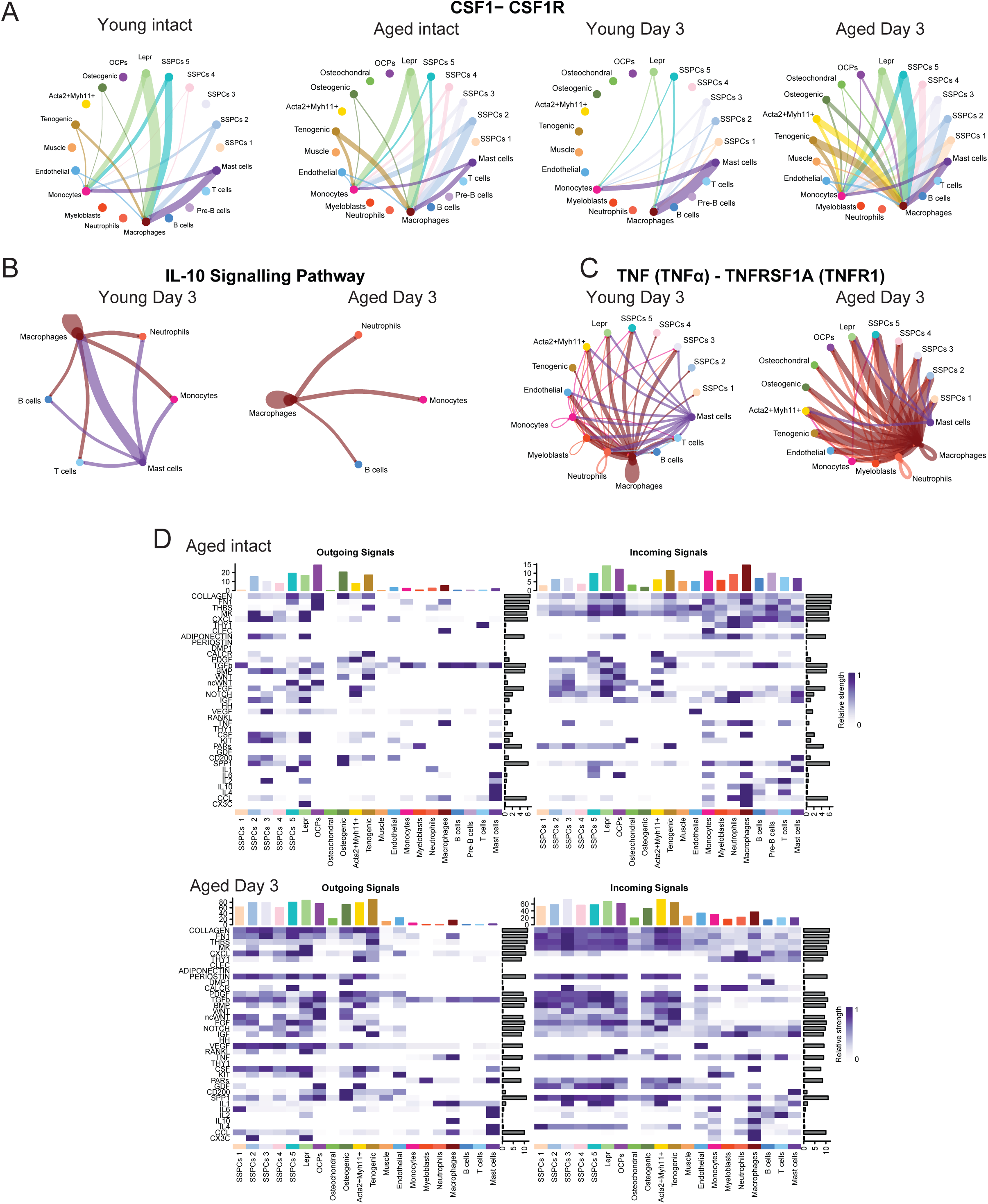
CellChat analysis of integrated CD45(-) and CD45(+) scRNA-seq datasets. **A)** Ligand-receptor interaction network for CSF1 to CSF1R in all cell types identified by scRNA-seq in merged CD45(-) and CD45(+), plus the individual clusters of the subsetted Prrx1+ pSSPCs, for each condition. **B)** Interactions for the IL-10 Signaling Pathways, and **C)** TNF-TNFR1 receptor-ligand interaction network three post-fracture, comparing young and old. The color of the line corresponds to the color of the dot representing the “sending cell” group within the interaction; a thicker line represents a stronger interaction. **D.)** Interactions between selected, enriched signaling pathways with outcoming and incoming signals for aged intact and three days post-fractured cells. The top bar plot shows the total signaling strength of a cell group, obtained by summarizing all signaling pathways displayed in the heatmap, and the right grey bar shows the total signaling strength of a pathway obtained by summarizing all cell groups displayed in the heatmap.

Our analysis further highlighted select communication patterns across different conditions: ’Aged Intact,’ ’Aged Day 3 Fracture,’ ’Young Intact,’ and ’Young Day 3 Fracture’ (Fig S8). In young intact periosteum, we found that Lepr+ cells were among the most active signal-sending cells, interacting primarily with other mesenchymal cell populations with strong CXCL signaling going to other immune cells. Notably, macrophages could be bioinformatically identified as major source of TNF with few cell-to-cell interactions at baseline. Following fracture, TNF signaling was strongly sourced from both macrophages and neutrophils, acting on many cell types (Fig S8A-D).

While interactions relating to collagen (COLLAGEN), fibulin (FN1), thrombospondins (THBS), and Midkine (MK) were the most enriched, we also assessed interactions related to skeletal regeneration, such as the BMP, WNT, and NOTCH pathways, and interactions related to inflammation, such as IL1 and TNF, among others (Fig 6D; Fig S8E-H). Our results demonstrated that signaling from CXCL primarily originated from the Lepr+ cells, which overlapped with CXCL12+ expressing cells, a key regulator of the bone marrow microenvironment. Additionally, we identified pathways that were unique to specific conditions, such as CX3CR signaling, which was primarily observed in aged mice post-fracture (Fig 6D; Fig S8E-H). Our CellChat analysis provides a comprehensive view of the cell-to-cell communication networks between the multiple cell types that drive fracture healing and identifies specific age-related alterations that may hinder bone regeneration.

## Discussion

Our study is a comprehensive analysis of the early periosteal response to fracture at the single-cell level in young and aged mice. It highlights significant age-related differences and interactions in mesenchymal and immune cell populations, contributing unique scRNA-seq datasets specifically relevant to understanding how aging affects periosteal cells during fracture healing. Notably, previous studies investing the periosteum at a single-cell level have relied on cells derived from ex vivo expanded periosteal cells (Julien et al., 2022), single nucleus periosteal cells (Perrin et al., 2024), or periosteal cells directly isolated from the fracture callus of young adult mice three days post-fracture, using a similar isolation method to this study (Novak et al., 2024).

Flow cytometry demonstrated an increased percentage of CD8+ T cells in the periosteum of aged mice three days post-fracture (Figure 4E). However, this increase in CD8+ T cells was not detected within the periosteal scRNA-seq dataset because T cell populations represent less than 5% of the cells (Fig 2C) Previous studies have shown that terminally differentiated CD8+ T cells and the adaptive immune system can broadly inhibit bone regeneration, which may contribute to age-induced alterations in fracture healing (Goodnough & Goodman, 2022; Reinke et al., 2013; Toben et al., 2011). While we noted a marked increase in CD8+ cytotoxic T cells three days post-fracture, Lopez et al. 2022 observed a significant increase only at 14 days post-fracture when assessing whole bone callus (39). In comparing our study to similar work assessing immune cell dynamics using flow cytometry, it is crucial to consider both the age of the mice and the skeletal compartment from which the cells were sourced. Previous studies have typically used cells from the whole tibia (Molitoris, Balu, et al., 2024), while we specifically focused on cells from the femoral periosteum. Additionally, prior studies have used cells from middle-aged mice (12 months old) (Lopez et al., 2022), whereas all aged mice in our study were 20-24 months.

Our scRNA-seq data comparing intact periosteum in young and aged male mice identified *Cxcl9* as one of the most differentially expressed genes, with significant upregulation in the aged pSSPC population. Others have found that increased CXCL9 is a risk-factor biomarker for osteoporotic hip fractures in males (Phan et al., 2022) and inflammatory aging (Sayed et al., 2021). Further, Cxcl9 expression is upregulated in whole bone from ovariectomized mice (Z. Liu et al., 2020) and was identified as the most upregulated gene in the long bone of aged male mice by bulk RNA-sequencing (Hebb et al., 2018). Although CXCL9 is known to regulate T-cell function via CXCR3, this receptor is minimally expressed in our CD45(+) scRNA-seq, possibly attributed to the few number of T cells detected, relative to other immune cell populations (Fig S4A-B). A prior study showed that CXCL9 impairs osteogenic processes though novel mechanisms by either direct inhibition on osteogenic culture differentiation or through antagonizing VEGF signaling (Huang et al., 2016) – an important pathway for proper osteoblastic activity during fracture repair (Hu & Olsen, 2016). However, VEGF-A (*Vegfa*) and its primary receptor VEGFR-2 (*Kdr*) are not differentially expressed in our datasets. Our CellChat data indicated that *Cxcl9* may be derived from, and act on, mesenchymal populations through the Atypical chemokine receptor 1 (*Ackr1*). Atypical chemokine receptors are thought to act as decoys for several chemokines, including CXCL9, with non-classical functions that have emerging importance in the immune system and in modulating hematopoietic stem cells (Comerford & McColl, 2024; Hur et al., 2016). While CXCL9 can also recruit CD8+ T cells, which are known to inhibit fracture healing (Reinke et al., 2013), our CellChat analysis did not detect a significant interaction between CXCL9 and CXCR3 on T cells, possibly because it was below the detection limit, as T cells made about 1% of the CD45+ cells. Through flow cytometry, we were able to more precisely quantity this population and show that aged mice had four times as many CD8+ T cells present three days after fracture in the periosteum compared with young mice (Fig 4F). These findings suggest that CXCL9 may adversely affect the periosteum and skeleton through multiple, potentially unknown mechanisms that warrant further investigation.

Distinct from bone marrow SSPCs, periosteal SSPCs (pSSPCs) are the primary contributors to the cartilaginous callus and new bone formation post-fracture (Jeffery et al., 2022; Li et al., 2022). Studies on heterochronic parabiosis between young and aged mice suggest that circulating factors may influence age-related bone repair (Baht et al., 2015). However, inherent alterations in skeletal stem cells, as demonstrated by in vivo renal transplantation assays of bona fide skeletal stem cells (CD51+THY1-6C3-CD105-CD200+) from aged mice, indicate intrinsic age-related defects in bone-forming capacity (Ambrosi et al., 2021). We observed a marked decline in the proliferative capacity of Prrx1+ pSSPCs in aged mice after fracture, evidenced by a significant reduction in cycling cells by EdU labeling and scRNA-seq (Fig 1B and 3F). The bioinformatically identified impaired proliferative response of Prrx1+ pSSPCs in aged mice coincided with increased cellular senescence markers, as indicated by the *SenMayo* gene signature. These data support the concept that aging reduces the availability and functional capacity of SSPCs essential for effective fracture healing (Fig 5K) (Ambrosi et al., 2021; Josephson et al., 2019). Whether this effect within periosteal SSPCs is cell-intrinsic, -extrinsic, or both remains to be determined.

In assessing transcriptional changes in pSSPCs post-fracture, we identified downregulation of chondrogenic gene sets, including *Col2a1* and *Acan*, in aged mice post-fracture, indicating reduced or delayed chondrogenic differentiation within periosteal cells as they age (Fig 5A-D). Aged mice further showed enrichment of immune response-related processes and gene sets (Fig 5A-D). These findings align with previous studies showing reduced chondrogenic gene expression and a smaller cartilaginous callus in aged mice (Clark et al., 2023), suggesting that early events in fracture healing significantly influence callus formation, setting the foundation for complete healing.

Integration of the datasets revealed distinct interaction patterns in periosteal cells derived from aged mice three days post-fracture, suggesting altered signaling dynamics in the aged periosteal microenvironment (Fig 6; Fig S8). While the microenvironment of the bone marrow has been a focus of other studies (Bandyopadhyay et al., 2024; Matsushita et al., 2020), interactions in the periosteum appear to have similar significance. Although the periosteum is traditionally viewed as a supportive structure, emerging studies suggest that the dura, which is a specialized periosteum of the inside of the skull (Rindone et al., 2021), has a reservoir of immune cells and channels between hematopoietic marrow in the interior of the calvarium (Cugurra et al., 2021; Smyth et al., 2024). Possibly, similar structures and resident immune cells contributing to a specialized microenvironment exist in long bones. This concept is especially relevant considering that periosteal-derived mediators of hematopoietic niches, such as Lepr+Cxcl12+ pSSPCs engaging in CXCL signaling, are present (Fig 3D; Fig S8). However, in our work, because a specific periosteal marker gene was not used to select for cells exclusively from the periosteum, such as Ctsk-linage mesenchymal cells (Debnath et al., 2018), it is possible that some of the Lepr+ cells represented in our single cell data set could have originated from the bone marrow or endosteal compartment.

One limitation of our study is that we only used male mice. Although both males and females experience age-related bone loss (Lu et al., 2022; Molitoris, Balu, et al., 2024), bone loss is significantly accelerated in females due to decreased estrogen receptor signaling following menopause or ovariectomy (OVX) in mice (Z. Liu et al., 2020). Clinically, osteoporosis and fragility fractures occur at a higher rate in females, with the biological impact of fractures further compounded by the adverse effects of estrogen withdrawal on fracture repair(Andrew et al., 2022; Walker & Shane, 2023). Consistent with our detection of upregulated Cxcl9 in aged periosteal cells, Liu et al. (Z. Liu et al., 2020) demonstrated that osteoblast-derived Cxcl9 in OVX mice contributes to the disruption of bone remodeling balance seen in postmenopausal osteoporosis. Future studies investigating the basal changes in the female periosteum caused by estrogen deprivation and aging could provide insights into how these processes compromise regenerative potential.

In summary, this study assessed how the periosteum and periosteal response to fracture is affected by aging in mice. Parallel scRNA-seq of CD45(+) and CD45(-) periosteal cells suggests multiple mechanisms are altered with age, potentially comprising bone healing in aged individuals. Changes in gene expression and the distribution of immune cell populations were corroborated by bulk RNA-seq and flow cytometry, respectively. Among CD45(-) cells, our findings highlight the reduced proliferative capacity of aged pSSPCs, their resistance to chondrogenic differentiation, and their senescence-associated gene expression signature. In the CD45(+) cell population, aged mice exhibited a marked increase in both CD4+ and CD8+ T cells post-fracture, whereas the fracture-induced increase in patrolling M2-like macrophages was diminished with age. These findings reinforce the concept that post-fracture, the aged periosteum creates an inflammatory microenvironment that induces a gene expression profile in CD45(+) cells consistent with a defensive immune response (Fig 5B; Fig 6A-B). Overall, our data highlight the role of immune cells and SSPCs intercellular communication and provide insight into how aging alters in fracture healing in ways that may compromise clinical outcomes.

## Materials and Methods

### Animal welfare and mice

This study was approved by the UConn Health Institutional Animal Care and Use Committee (IACUC). All animals were housed in ventilated cages in a temperature and humidity-controlled environment under a 12-hour light cycle. This study used strain-matched C57BL/6 male mice. All young mice were purchased from Charles River Laboratories and were approximately 3-months of age at the time of experiments. All aged mice were supplied by the National Institute on Aging’s Aged Rodent Colonies at 18 months of age with experiments conducted when the mice were between 20-24 months of age.

### Surgical procedure

Closed pin stabilized bilateral femoral fractures were generated in young and aged mice as previously described (Doherty et al., 2023; Manigrasso & O’Connor, 2004). Briefly, pin placement was confirmed using an X-ray system (Faxitron or Kubtec), and the femur was fractured using an Einhorn three-point drop-weight device and evaluated using X-ray. Buprenorphine (0.1 mg/kg body weight) was administered by subcutaneous injection before the procedure and twice a day for three days following the fracture for pain management. In some experiments, mice were injected with buprenorphine extended-release injectable suspension (Ethiqa XR; Fidelis Animal Health, Inc) 1.3 mg/ml once after the surgery. Samples excluded from the study analysis included multiple (comminuted) fractures, bent pins, and fracture placements that were not mid-diaphyseal. Mice were euthanized by CO_2_ asphyxiation on post-fracture operation day three to isolate bones.

### Isolation of cells for scRNA-sequencing, RNA-sequencing, and flow cytometric analysis

Periosteal cells were isolated from intact femurs as previously described (Scanlon et al., 2017). For fractured femurs, after removing the muscle, the periosteum at the fracture ends was visualized using a dissecting microscope and gently scraped into sterile PBS and digested enzymatically (0.05% collagenase P and 0.2% hyaluronidase in PBS) for 45 minutes at 37 °C in an orbital shaker. Cells were resuspended in the staining media (0.1M HEPES and 2% heat-inactivated FBS in Hanks Balanced Salt Solution (HBSS) and passed through an 18-gauge needle five times to break clumps. After filtering the suspension through a 70 μm mesh strainer to remove debris, cell suspensions were centrifuged at 300 x g for 5 minutes at 4°C. The cell pellet was resuspended in the staining media and processed for flow cytometry, FACS isolation, or magnetic bead separation. Blood was drawn by cardiac puncture using an EDTA-coated insulin syringe, then added to a tube containing 0.2 ml of EDTA (10 mM) solution. After gentle mixing, 1 ml ACK buffer (Sigma) was added to lyse red blood cells. After 5 minutes of incubation at room temperature, PBS (9 ml) was added, and cells were centrifuged again at 300 x g for 5 minutes at 4°C. This process was repeated twice, and after the final wash, cells were suspended in the staining media.

### Immunophenotyping by Flow Cytometry

Periosteal cells from intact and fractured femurs were collected as described above. Each mouse was treated as one replicate for flow cytometric analysis of immune cells. After isolation, cells (1×10^6^) were incubated with the antibodies indicated in Supplementary Table 2 and processed for flow cytometry as previously described (Doherty et al., 2023). Dead cells were excluded using 4, 6-diamidino-2-phenylindole (DAPI) staining. Flow cytometry was performed on a BD FACSymphony A5 and analyzed in FlowJo (v10) (BD). All experiments included unstained bone marrow and periosteal cells for establishing gates. Fluorochrome beads were used as single-color controls. Unstained, single-stained, and fluorescence minus one (FMO) controls were used in establishing initial gates and compensation using cells isolated from the bone marrow.

### Single-cell RNA-sequencing

Periosteal cells from intact and fractured femurs were collected as described above. For scRNA-sequencing, periosteal cells from 5-7 mice were pooled together for each time point and condition. To enrich the CD45(-) cell population for scRNA sequencing, cells were stained with the appropriate antibodies described above to sort for live Ter119(-)CD45(+) or Ter119(-)CD45(-) cells using a FACSAria II (BD), which were used as input for single-cell sequencing. The Single Cell Biology Services (SCBL) at The Jackson Laboratory for Genomic Medicine (Farmington, CT) performed FAC-sorting and 10x Chromium droplet-based single-cell library preparation and sequencing for 5’ Gene Expression using 6000 live cells as input, which were sequenced to a target read depth of 125K/cell for the CD45(-) cell datasets and 100K/cell for the CD45(+) cell datasets. We subsequently used FASTQ files and Cell Ranger (v7.0.1) for gene counting with the force cell function set to 6000. Filtered feature barcode matrix files were used for downstream clustering and analysis using Seurat-based pipelines (v4) and reference vignettes from its creators (Satija Lab) (Hao et al., 2021) in RStudio (v4). To remove low-quality cells, we filtered cells based on *nFeature_RNA* (>200 or <10,000), mitochondrial content (<30% *mt-* related genes), and hemoglobin-related genes (<2% *Hbb-* related genes). To integrate multiple datasets together, we used the SCTransform (v2) normalization method (Hafemeister & Satija, 2019). Trajectory analysis and CellChat analysis were based on vignettes and publicly available GitHub scripts, referenced by Cao et al. 2019, Jin et al. 2021, and Bandyopadhyay et al. 2024 (Bandyopadhyay et al., 2024; Cao et al., 2019; Jin et al., 2021).

### Bulk mRNA sequencing and analysis

Periosteal cells from intact and fractured femurs were collected as described above. In this experiment, 2-3 mice were pooled together to create one replicate. To isolate cells for bulk RNA-sequencing, cells were incubated with anti-mouse CD45 phycoerythrin (PE) tethered antibodies (1:200) for 30 minutes on ice. After washing twice with the staining media, cells were incubated with anti-PE MicroBeads, and CD45(-) cells were eluded using MACS separation Columns (Miltenyi Biotec) according to the manufacturer’s protocol. RNA was isolated from freshly isolated, magnetic sorted, CD45(-) cells using a Trizol (Invitrogen) extraction method. Total RNA was quantified, and only samples with RNA integrity values above 8.0 were included for library preparation. Illumina transcriptome library preparation and sequencing total RNA samples were prepared for mRNA-sequencing using the Illumina Stranded mRNA Ligation Sample Preparation kit following the manufacturer’s protocol (Illumina). Libraries were validated for length and adapter dimer removal using the Agilent TapeStation 4200 D1000 High Sensitivity assay (Agilent Technologies), then quantified and normalized using the dsDNA High Sensitivity Assay for Qubit 3.0 (Life Technologies). Sample libraries were prepared for Illumina sequencing by denaturing and diluting the libraries per the manufacturer’s protocol (Illumina). All samples were pooled into one sequencing pool, equally normalized, and run as one sample pool across the Illumina NovaSeq 6000 using version 1.5 chemistry. A target read depth of 50 M reads was achieved per sample paired-ended end 100 bp reads. Raw reads were trimmed with *fastp* (v0.23.0), with low-quality sequences removed, and trimmed reads mapped to the *Mus musculus* genome (GRCm39) with HISAT2 (v2.1.1)(Kim et al., 2019). SAM files were then converted into BAM format using *samtools* (version 1.12). Counts were generated against the features with 100 HTSeq-count. Differential expression of genes between conditions was evaluated using DESeq2(Love et al., 2014). When analyzing our bulk RNA-seq datasets for differential expressional analysis, some genes classified as pseudogenes, micro RNAs (miRNAs), undefined, long non-coding RNAs (lncRNAs), ribosomal, canonical genes associated with hematopoietic cells and progenitor, such as hemoglobin, and mitochondrial genes were removed. Gene Set Enrichment Analysis (GSEA) was performed using the Gene Ontology Biological Process datasets with the *fgsea* package (Korotkevich et al., 2016).

### Histology

To assess cell proliferation, mice were injected with 5-ethynyl-2′-deoxyuridine (EdU) (10 mM) in normal saline 24 h before sacrifice. Fractured and intact femur sections obtained from separate cohorts of young and aged mice were stained for EdU labeling using the Click-iT® EdU Alexa Fluor® 488 Imaging Kit (Life Technologies/Thermo Fisher Scientific) according to the manufacturer’s instructions. Sections were rinsed in PBS, counterstained with DAPI to visualize nuclei, and mounted for fluorescent imaging. Fluorescent images were visualized using an Eclipse 50i microscope (Nikon) or an Axioscan slide scanner (ZEISS). Within the ZEN (blue edition) software (ZEISS), a region of interest was applied to each side of each fracture callus. These images were analyzed in ImageJ (NIH) by first thresholding for EdU-labelled cells, then transformed using the watershed function, and automatically counted. The exact process was performed for DAPI-labeled, yielding the proliferating fraction. Alkaline Phosphatase (ALP) activity was evaluated using a modification of the Leukocyte ALP Kit (Sigma-Aldrich) and calculated using ImageJ, where the mean intensity of the ALP signal was normalized to a given region of interest area.

### Data analyses

Statistical analyses were performed using RStudio (v4.3.1) and GraphPad Prism (v10), with figures assembled in Adobe Illustrator (v28.1). Data are presented as mean ± SD. P-values < 0.05 were considered statistically significant.

## Ethics Statement

All authors have no disclosures pertaining to this work.

## Data availability

The data that support the findings of this study are openly available in the Gene Expression Omnibus at https://www.ncbi.nlm.nih.gov/geo/query/acc.cgi?acc=GSE280914 reference number GSE280914.

## Abbreviations

SSPC: skeletal stem and progenitor cells
scRNA-seq: single-cell RNA-sequencing

## Acknowledgments

We gratefully acknowledge the contribution of the Single Cell Biology Services (SCBL) at The Jackson Laboratory for expert assistance with the work described herein. These shared services are supported in part by the JAX Cancer Center (NIH/NCI P30 CA034196). We would like to thank Dr. Vijender Singh and the UConn Center for Genome Innovation for bulk RNA-sequencing and computational support, and Sangit Karki for the technical assistance. BioRender was used to construct graphics and illustrations. This study was supported by grant funding to AS and AMD (NIH/NIA R21-AG071047), IK (NIH/NIAMS R01-AR055607), and JSK (NIH/NIDCR T90-DE033006).

**JSK**: conceptualization and writing – original draft, editing, and final version; experimental investigation; bioinformatic analysis. **MW**: experimental investigation. **AK:** bioinformatic insight. **SN:** experimentation investigation. **IK:** experimentation investigation and intellectual insight. **AD:** conceptualization; intellectual insight; funding, revision, and editing. **AS**: project supervision; funding; conceptualization and writing – revision and editing. All authors read and approved the final version of the manuscript

---

**Supplemental Figure 1.**
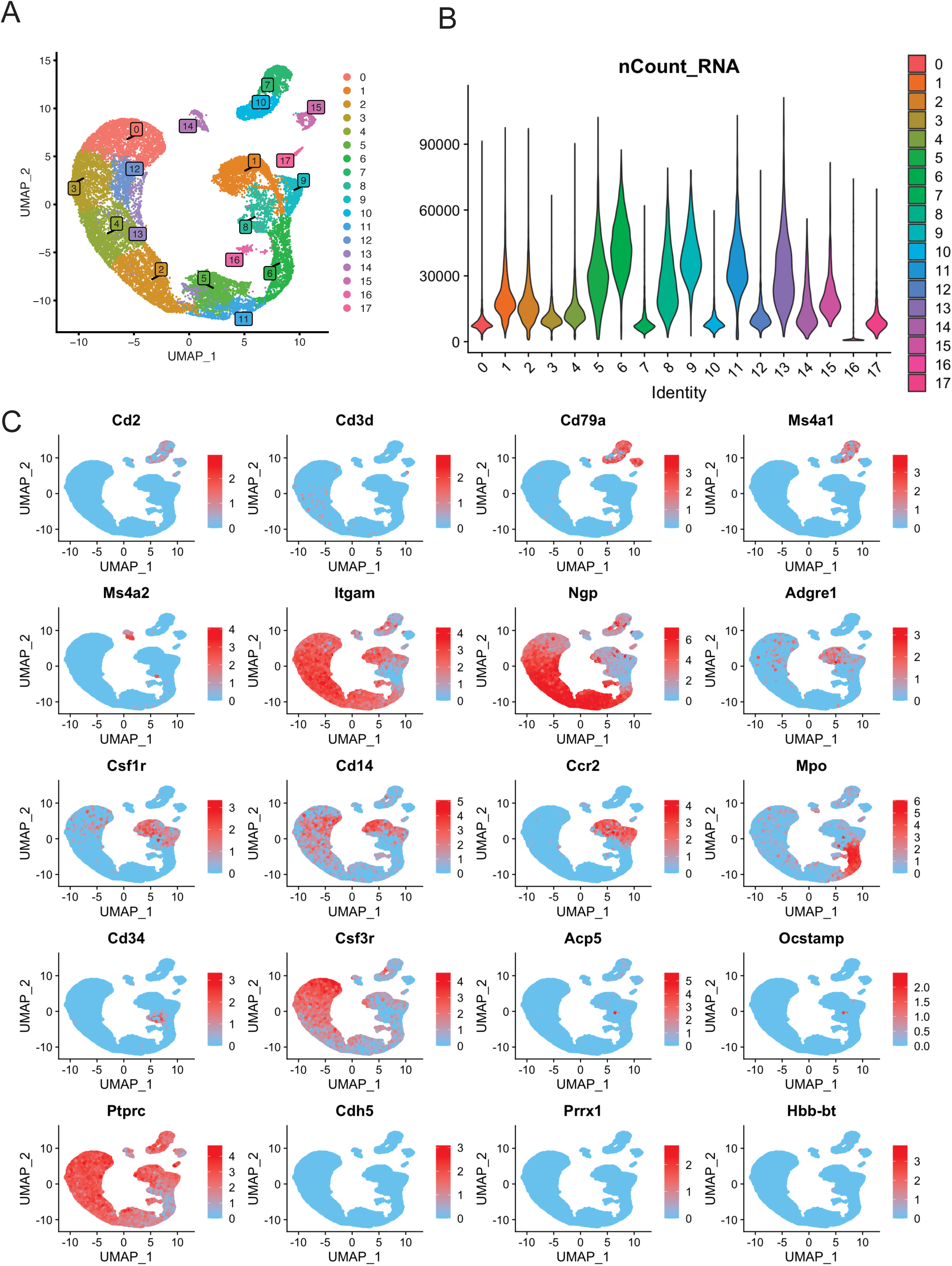
Cluster identities of CD45(+) scRNA-sequencing data. **A)** Initial clustering of the merged dataset before assigning identities. **B)** Plot of *nCount_RNA* (RNA content in each cell) for each cluster, used to identify low RNA-content cells, such as neutrophils. **C)** Feature plots and the *FindMarkers* function in Seurat were used to identify each cluster’s cell type based on several markers. Neutrophils, marked by *Ngp*, *Itgam*, and *Csf3r*, were identified in Clusters 0, 2, 3, 4, 12, and 13, which were subsequently merged. Macrophages, marked by *Cd14*, were found in Cluster 1, while monocytes, marked by *Adgre1*, were identified in Clusters 8 and 9. Cluster 14 was enriched with mast cell markers, such as *Ms4a2*. Lymphocyte clusters included B cells (marked by *Cd79a* and *Ms4a1*/CD20) in Cluster 10, and pre-B cells (marked by *Cd79a*) in Cluster 15, which displayed more immature markers. Cluster 17, enriched for *Cd2* and *Cd3d*, was annotated as T cells. Clusters 5, 6, and 11 showed markers of myeloblasts. Cluster 16, which lacked discernible enrichment for specific cell-type markers, was annotated as ’non-specific’.

**Supplemental Figure 2.**
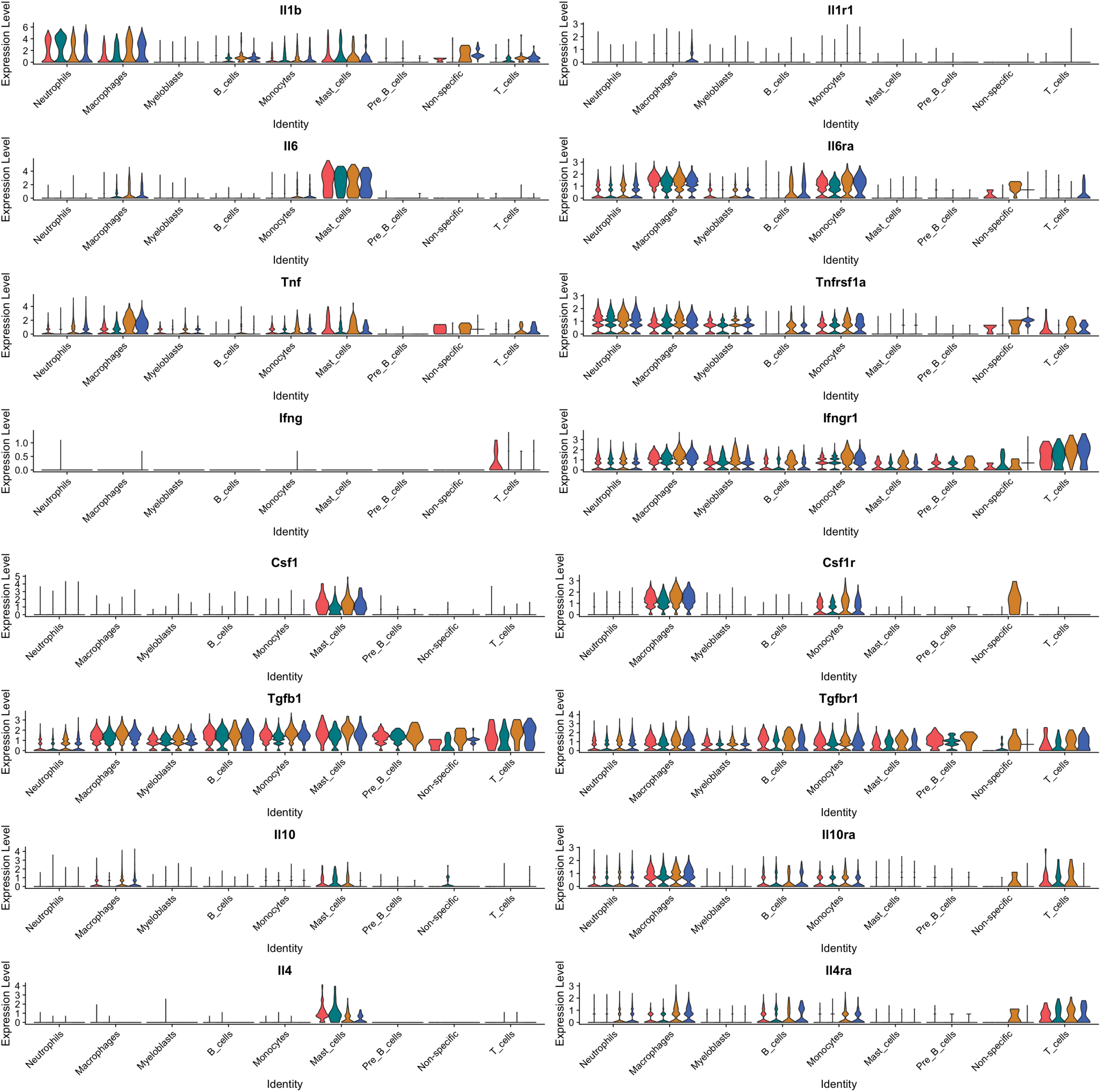
Violin plot of select cytokines/receptors from CD45(+) scRNA-seq datasets. The sample order for each gene is young intact (red), aged intact (teal), Young three days fracture (orange), and aged three days post-fracture (blue).

**Supplemental Figure 3.**
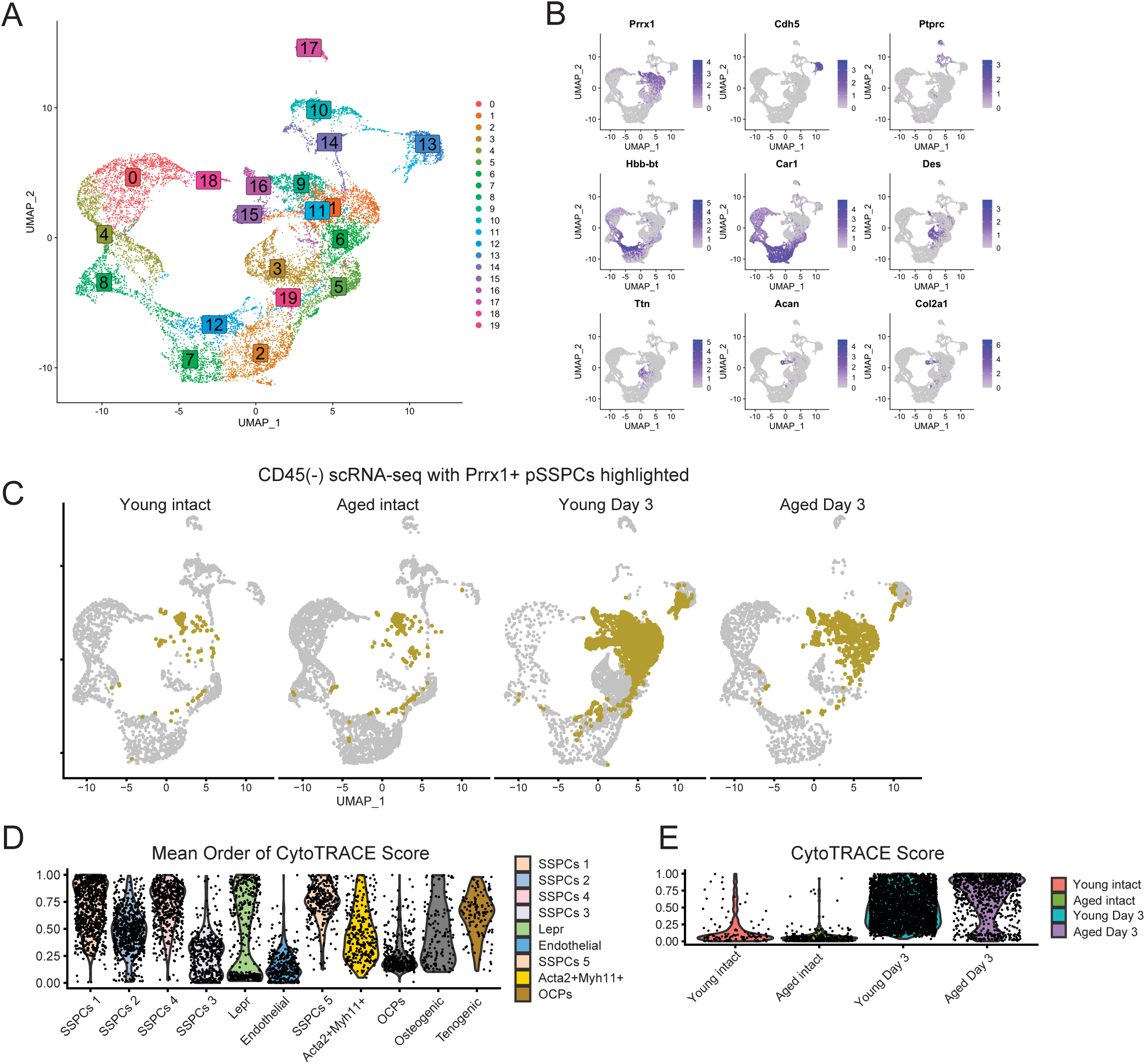
Data used to develop initial cluster identities in the CD45(-) scRNA-sequencing data. **A)** Initial clustering of the merged dataset before assigning identities **B)** Feature plots used to identify major cell types associated with each cluster. This was used in combination with the *FindMarkers* function in Seurat to identify each cluster in Fig 3A. **C)** The Prrx1+ pSSPC cell clusters highlighted in the complete CD45(-) datasets separated by group. **D)** Violin plot of the mean CytoTRACE score for each subsetted and reclustered Prrx1+ pSSPC group, which are arranged in descending order of mean CytoTRACE Score. **E)** Violin plot of the aggregate CytoTRACE score for each subsetted and reclustered Prrx1+ pSSPC group analyzed together.

**Supplemental Figure 4.**
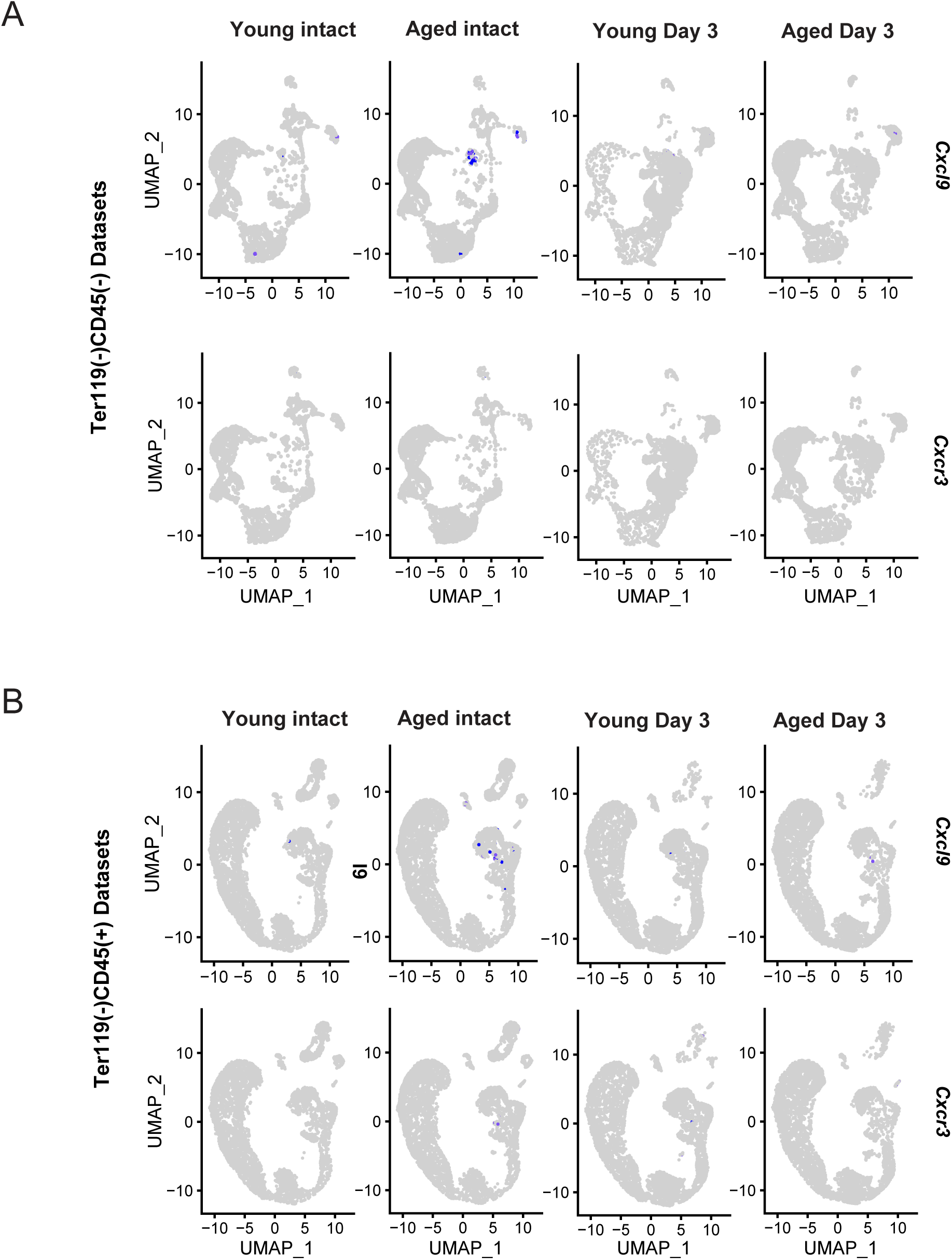
C*x*cl9 and *Cxcr3* expression. A) Feature plots based on the CD45(-) datasets. B) Feature plots based on the CD45(+) datasets.

**Supplemental Figure 5.**
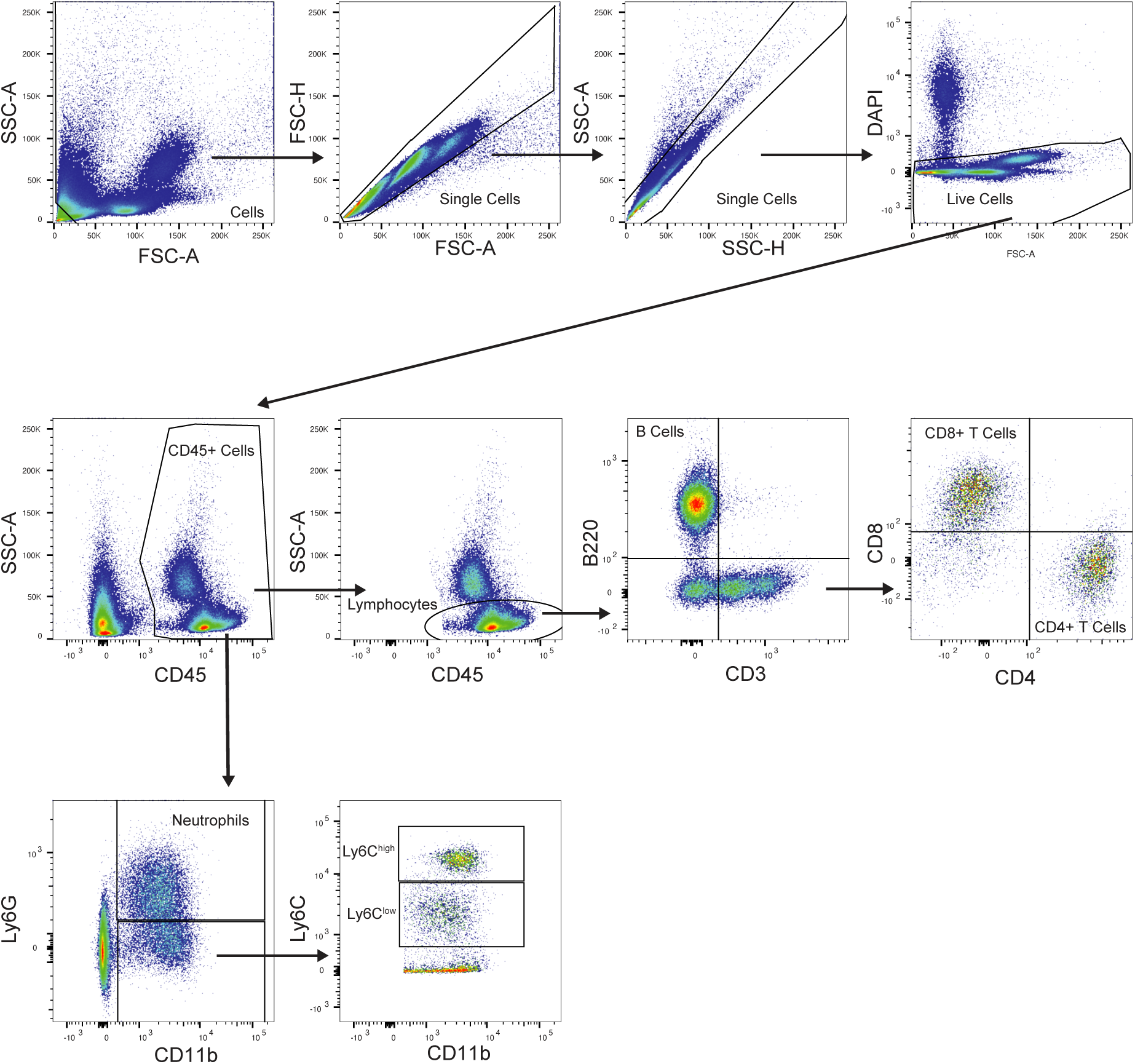
A representative gating strategy for flow cytometry is used to analyze immune cells from the periosteum, bone marrow, and peripheral blood. The gate for the FSC-A by SSC-A goes outside the field of view and includes all except in the lower left corner.

**Supplemental Figure 6.**
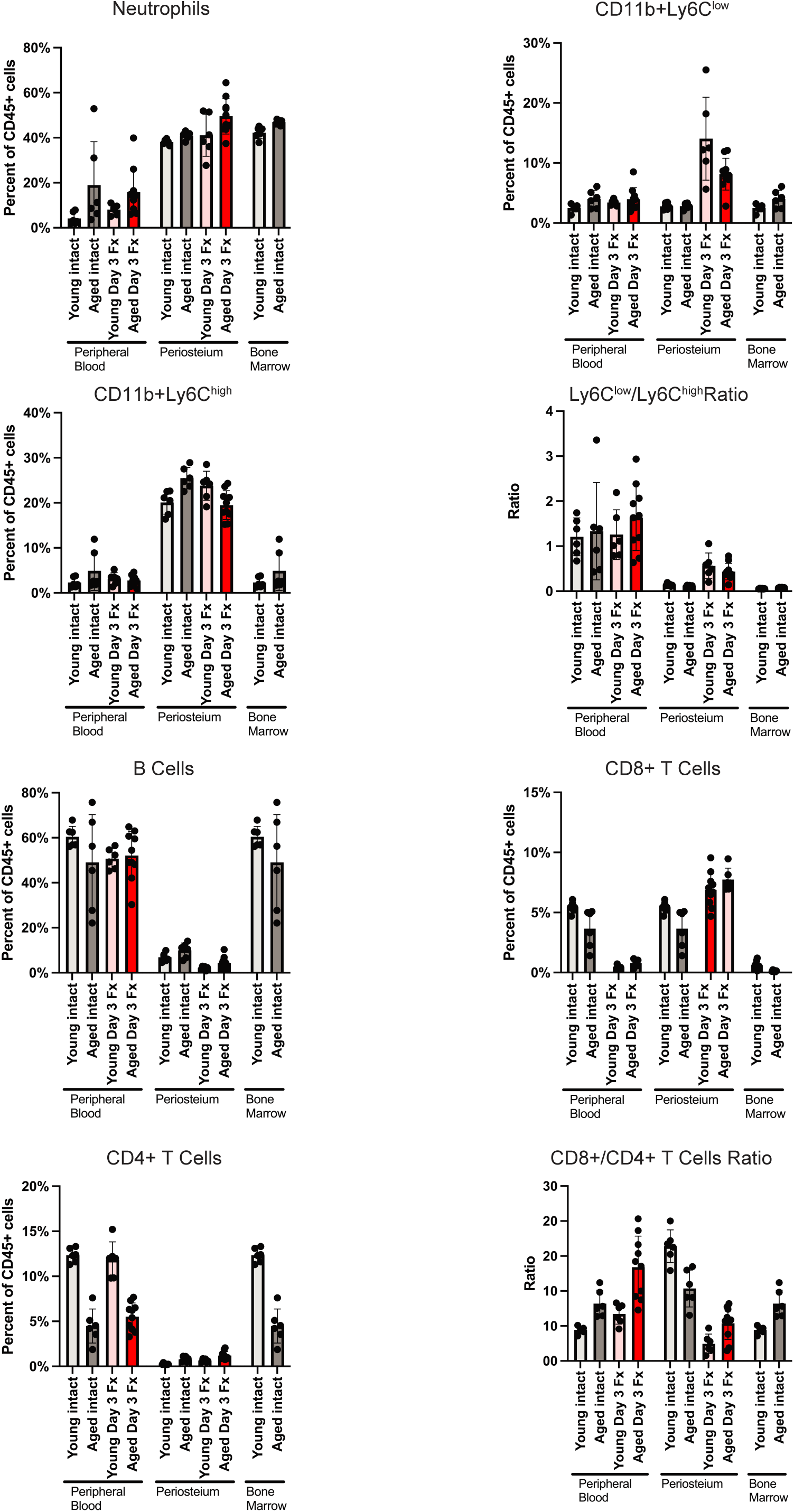
Comparing proportions of immune cells in the periosteum, whole blood, and bone marrow by flow cytometry, reported as a percentage of total CD45(+) cells. Samples of bone marrow were from intact femurs, as the use of an intramedullary pin in the femoral fracture model and soft callus at three days post-fracture prevents bone marrow flush. Each dot on the graph represents an individual animal as one biological replicate. N = 6-10/group.

**Supplemental Figure 7.**
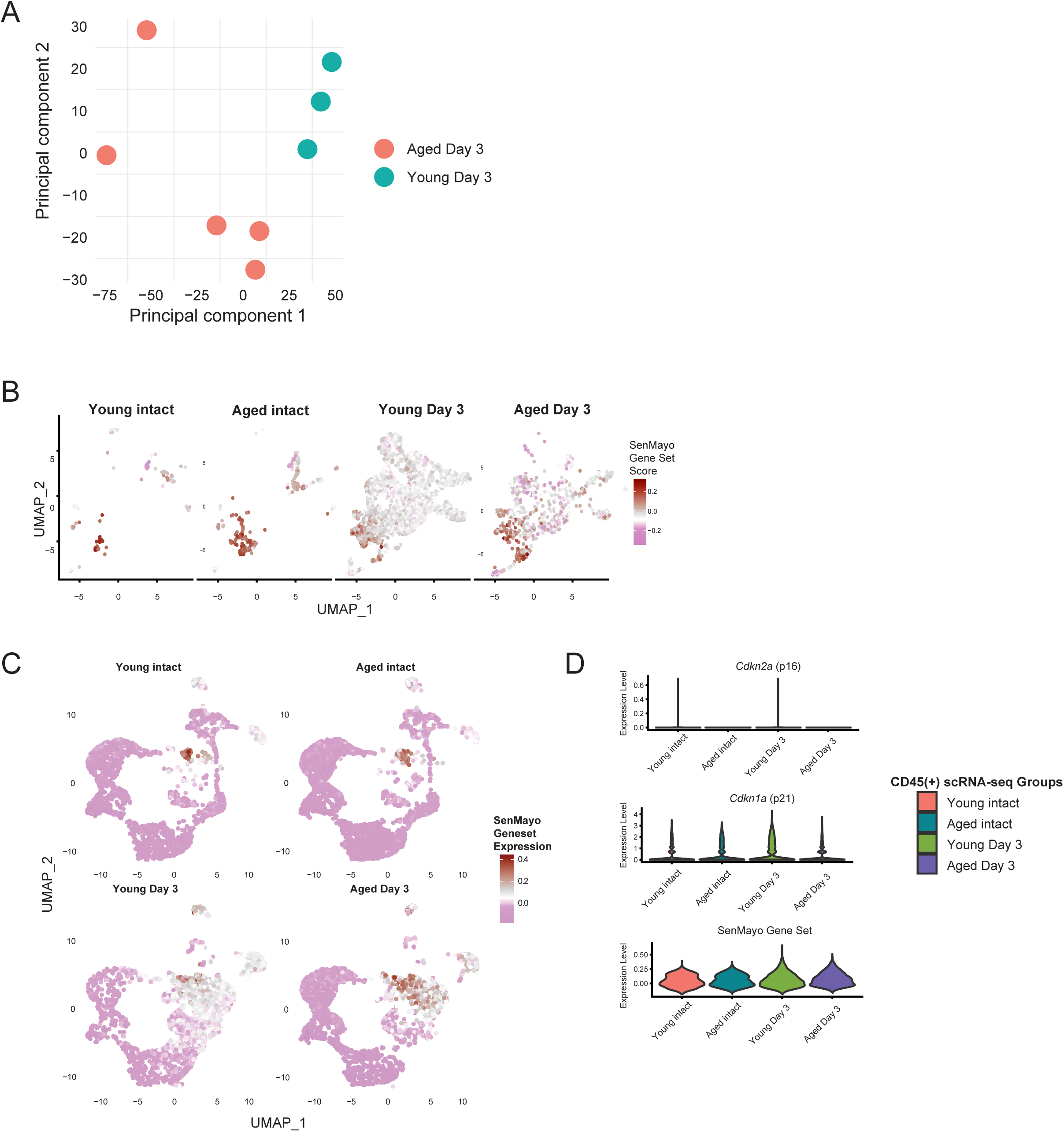
Analysis of periosteum by bulk RNA-sequencing **A)** PCA-plot of bulk RNA-seq samples isolated from CD45(-) periosteal cells from young and aged mice three post-fracture; each sample is composed of cells pooled from 2-3 mice. N=3 for young; N=5 for aged. Assessing periosteal cell senescence burden by scRNA-seq. **B)** Feature plot of the *SenMayo Gene Set Score* used to assess cellular senescence for each of the subsetted and reclustered Prrx1+ pSSPCs that was also **C)** applied to all cells of each condition of the CD45(-) scRNA-seq datasets. **D)** Analysis for p16, p21, and the SenMayo gene set applied to all cells of each group of the CD45(+) scRNA-seq datasets, which contained most of the immune cells.

**Supplemental Figure 8.**
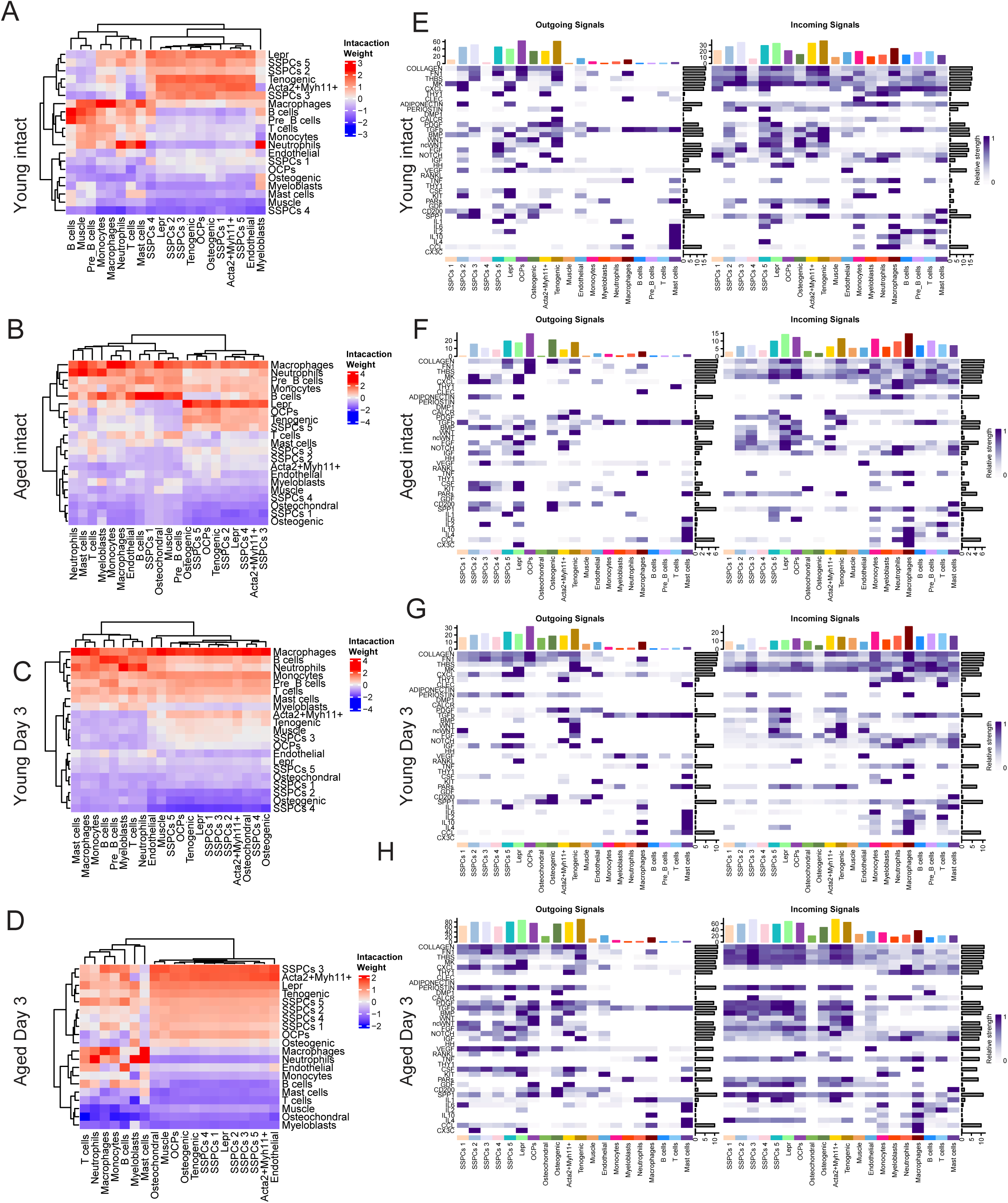
Cell-to-cell interaction networks **A-D)** Dendrogram heatmaps for interactions between cell types where the receiver cells are listed on the x-axis and the sender cells are listed on the y-axis, for each experimental group. **E-H)** Interactions between enriched signaling pathways, with outcoming and incoming signals for each of our groups. The top bar plot shows the total signaling strength of a cell group obtained by summarizing all signaling pathways displayed in the heatmap, and the right grey bar shows the total signaling strength of a pathway obtained by summarizing all cell groups displayed in the heatmap.

**Supplemental Table 1:**
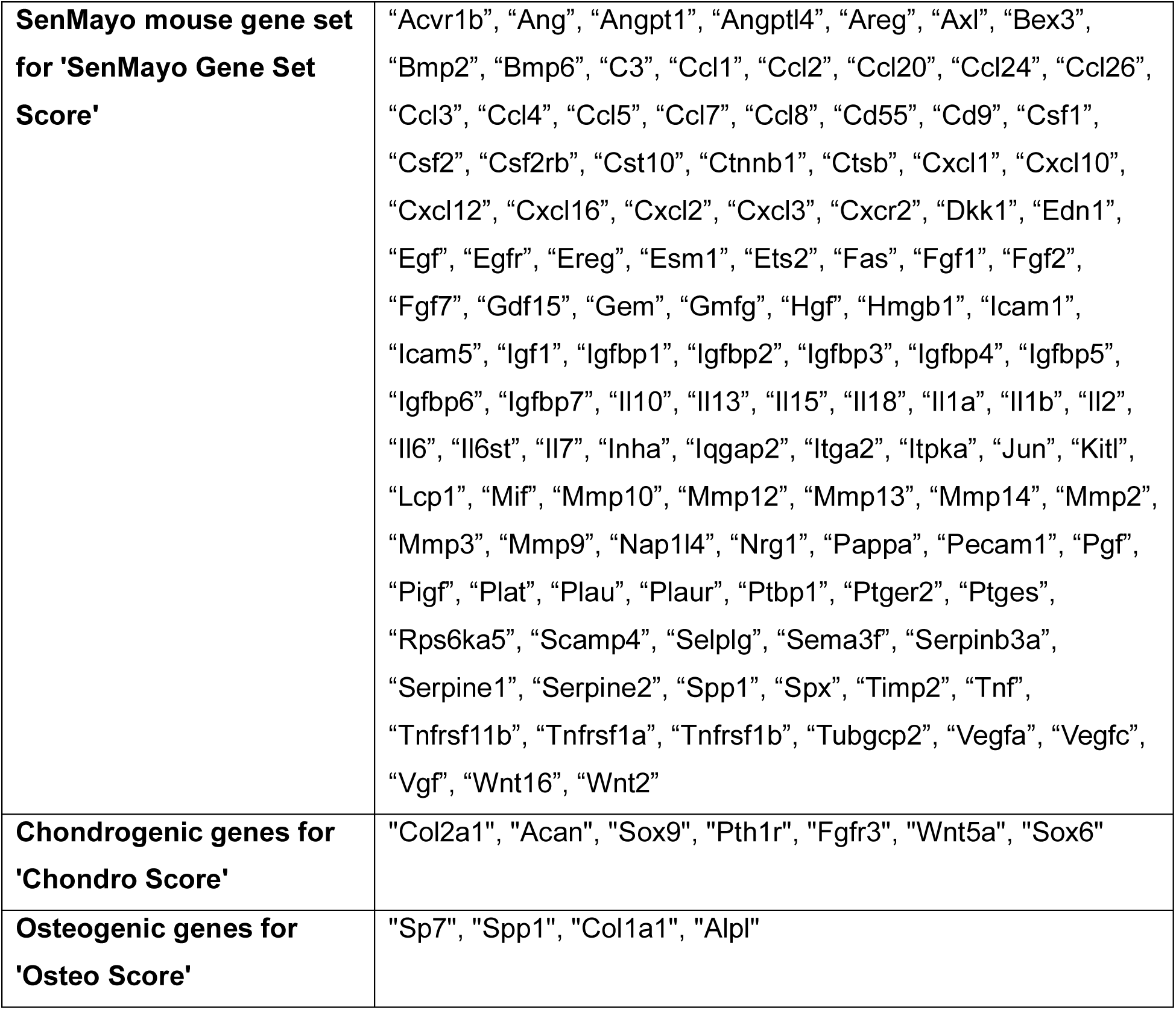
List of genes used for SenMayo-, Osteo-, and Chondro-score analyses in scRNA-seq analyses using the Seurat-based *AddModuleScore* function. The SenMayo gene list is based on Saul et al. 2022, and the Chondrogenic gene list was based on Julien et al. 2022.

**Supplementary Table 2.** Table listing antibodies and dilutions.

| Reagent Type | Designation | Source or reference | Identifiers | Additional information |
| --- | --- | --- | --- | --- |
| Antibody | Anti-Mouse TER-119 PE | eBioscience | 12-5921-82 | Flow (1:400) |
| Antibody | Anti-Mouse CD31 PE | eBioscience | 12-0311-82 | Flow (1:400) |
| Antibody | BD OptiBuild™ BUV737 Rat Anti-Mouse CD45 | BD Biosciences | 748371 | Flow (1:400) |
| Antibody | PE/Cyanine7 anti-mouse Ly-6C Antibody | BioLegend | 128017 | Flow (1:400) |
| Antibody | PerCP/Cyanine5.5 anti-mouse Ly-6G Antibody | BioLegend | 127615 | Flow (1:400) |
| Antibody | FITC anti-mouse CD3ε Antibody | BioLegend | 100306 | Flow (1:200) |
| Antibody | Anti-Mouse CD4 APC-eFlour® 780 | eBioscience | 47-0042-82 | Flow (1:400) |
| Antibody | Anti-Mouse CD8a Alexa Fluor® 700 | eBioscience | 56-0081-82 | Flow (1:400) |
| Antibody | APC anti-mouse/human CD45R/B220 Antibody | BioLegend | 103212 | Flow (1:400) |
| Antibody | CD11b Monoclonal Antibody (M1/70), eFluor™ 450, eBioscience™ | Invitrogen | 48-0112-82 | Flow (1:400) |
| Nucleic Acid Stain | DAPI | Fisher Scientific | P | Flow (1:1000) 3 μM final, 1.5 mM stock solution |

